# Endogenous Opioid Signaling Regulates Proliferation of Spinal Cord Ependymal Cells

**DOI:** 10.1101/2023.09.07.556726

**Authors:** Wendy W.S. Yue, Kouki K. Touhara, Kenichi Toma, Xin Duan, David Julius

## Abstract

After injury, mammalian spinal cords develop scars to seal off the damaged area and prevent further injury. However, excessive scarring can hinder neural regeneration and functional recovery (1, 2). These competing actions underscore the importance of developing therapeutic strategies to dynamically modulate the extent of scar formation. Previous research on scar formation has primarily focused on the role of astrocytes, but recent evidence suggests that ependymal cells also participate. Ependymal cells normally form the epithelial layer encasing the central canal, but they undergo massive proliferation and differentiation into astroglia following certain types of injury, becoming a core component of scars (3–7). However, the mechanisms regulating ependymal proliferation *in vivo* in both healthy and injured conditions remain unclear. Here, we uncover an intercellular kappa (κ) opioid signaling pathway that controls endogenous ependymal proliferation. Specifically, we detect expression of the κ opioid receptor, OPRK1, in a functionally under-characterized cell type called cerebrospinal fluid-contacting neurons (CSF-cNs). We also discover a neighboring cell population that express the cognate ligand, prodynorphin (PDYN). Importantly, OPRK1 activation excites CSF-cNs, and systemic administration of a κ antagonist enhances ependymal proliferation in uninjured spinal cords in a CSF-cN-dependent manner. Moreover, injecting a κ agonist reduces the proliferation induced by dorsal hemisection. Altogether, our data suggest a regulatory mechanism whereby PDYN^+^ cells tonically release κ opioids to stimulate CSF-cNs, which in turn suppress ependymal proliferation. This endogenous pathway provides a mechanistic basis for the potential use of κ opiates in modulating scar formation and treating spinal cord injuries.

---

Cerebrospinal fluid (CSF) flows through the central canal of the spinal cord, transporting metabolites into and away from the nervous system. The canal, which traverses the length of the cord along its midline, is lined by a layer of ciliated epithelial cells that retain proliferative capacity in adulthood. Following spinal cord injury, these ependymal cells divide and differentiate into astrocytes, thereby contributing to scar formation that limits secondary damage (3–7). Currently, it remains unknown how ependymal cell proliferation is regulated endogenously under healthy or pathological conditions, although the inhibitory neurotransmitter GABA has been suggested to be an instructive signal (8).

One potential source of GABA is a group of morphologically distinct cells called CSF-contacting neurons (CSF-cNs) that are highly conserved across vertebrate species. In mice, the somata of these cells are scattered within the ependyma, from which they extend a thin projection into the central canal, terminating in a bulbous enlargement that contacts the CSF (9). This characteristic morphology is reminiscent of olfactory sensory neurons and, indeed, CSF-cNs have been suggested to possess sensory functions, such as detecting changes in pH or osmolarity within the CSF (10, 11) or, in zebrafish, detecting the presence of meningitis bacteria (12). Zebrafish CSF-cNs have also been proposed to mechanically detect spinal curvature and provide motor feedback for controlling tail movement (13, 14). Nonetheless, whether mammalian counterparts have preserved these same functions or acquired new physiological roles is unclear.

In this study, we focus on the role of CSF-cNs in injury by asking whether and how they regulate ependymal cell proliferation under normal or pathological conditions. We find that CSF-cNs are constitutively activated by kappa (κ) opioids, which are produced by a neighboring cell type within the ependyma. Paracrine opioid signaling tonically suppresses ependymal cell proliferation. After partial spinal cord transection, we show that this suppression by CSF-cNs is relieved, revealing a mechanism for how ependymal cell proliferation is dramatically upregulated upon injury.

## Genetic labelling of CSF-cNs

The dearth of knowledge regarding mammalian CSF-cNs reflects, in large part, the lack of specific genetic access to these cells. Mouse CSF-cNs express the PKD2L1 ion channel, and thus most previous studies have used a *Tg(PKD2L1-Cre)* mouse line (10) to fluorescently label these cells. This method, in conjunction with the characteristic shape and location of CSF-cNs, has permitted their identification for electrophysiological recording and imaging (10, 11, 15–17). However, we found that this transgenic line also marks many other cells in the spinal cord (Supplementary Fig. 1a), making it difficult to trace and manipulate CSF-cNs selectively. This problem does not appear to be the result of transient *Pkd2l1* expression during development since non-CSF-cNs were labelled when a fluorescent reporter was introduced after adulthood using an AAV vector (Supplementary Fig. 1b). To bypass this issue, we asked whether particular AAV serotypes would show specific tropism for CSF-cNs when introduced into the CSF via the brain ventricles (i.e., intracerebroventricular (i.c.v.) injection, see Methods), which are connected to the central canal. Remarkably, AAV serotype 2/2 infected a set of cells around the central canal in a highly specific manner, even without the need to utilize a genetic Cre driver line. The AAV2-labelled cells express the CSF-cN marker, PKD2L1 (Fig. 1a) and demonstrate the stereotypical morphology of CSF-cNs (Fig. 1b). Using this method, we delivered an alkaline phosphatase (PLAP) reporter into CSF-cNs to reveal their detailed morphology. We observed that CSF-cNs send projections to the ventral white matter, in addition to the central canal (Fig. 1c). The same projection pattern was observed for CSF-cNs at all levels of the spinal cord up to the brainstem (Supplementary Fig. 1c). In sagittal sections, these ventral projections were seen to form a bundle that ran parallel to the central canal (Supplementary Fig. 1d). By titrating the concentration of injected AAV to achieve sparse labelling, the long projection of a CSF-cN could be seen to first extend ventrally from the soma, then turn 90° and extend rostrally for at least 400 μm (Supplementary Fig. 1e and g). These morphological features are comparable to those described in zebrafish (18) and, more recently, in mice using a similar but independently developed AAV injection method employing AAV2/1, which is reportedly less specific and preferentially labels CSF-cNs that are located ventrally in the ependyma (19).

**Figure 1.**
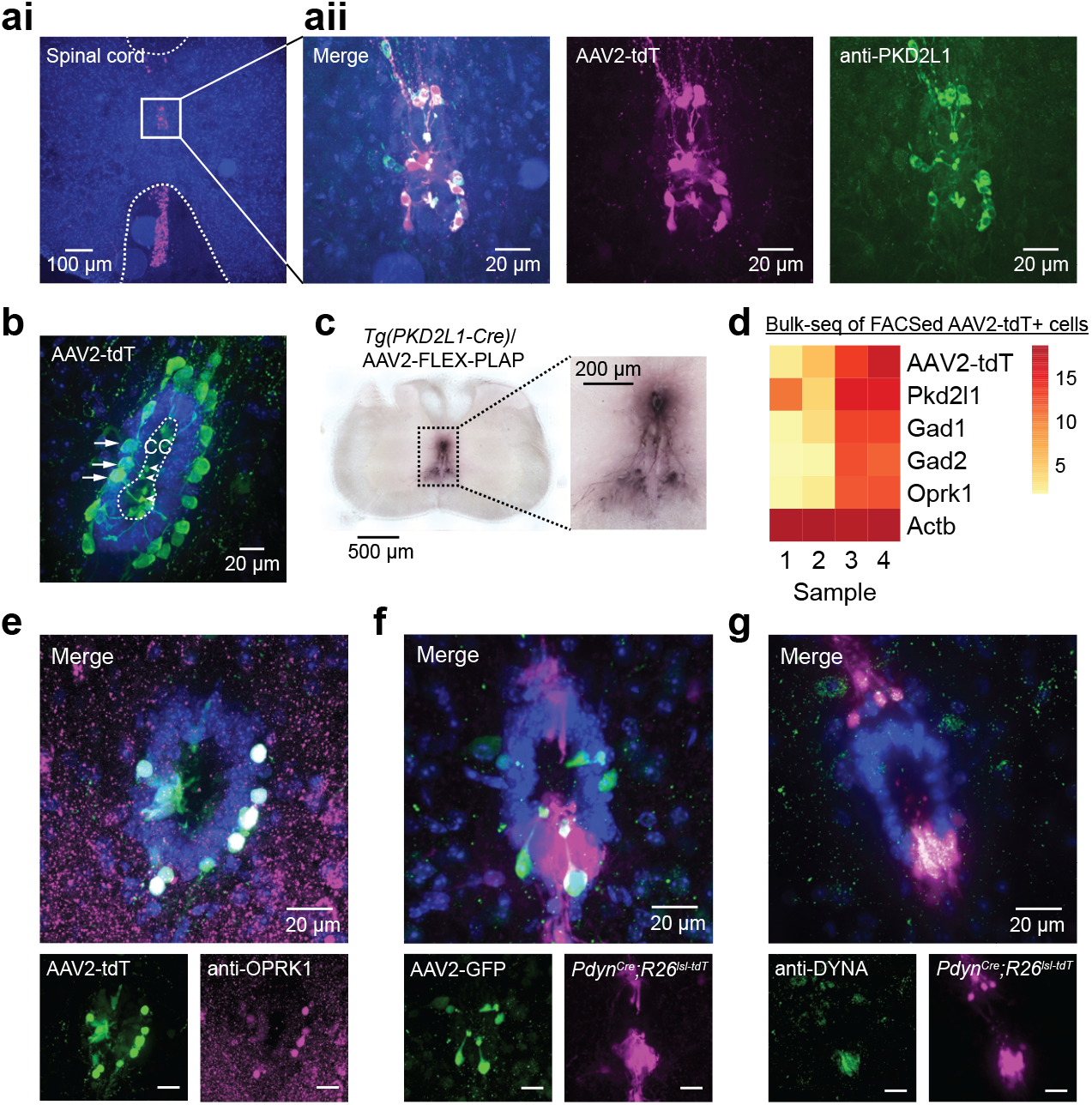
κ-opioid receptor and ligand are expressed in the ependymal region of the mouse spinal cord. **a**, Genetic labelling of CSF-cNs by intracerebrospinal AAV injection. **ai**, A representative coronal spinal cord section from a mouse with AAV2/2-tdTomato injected to its right lateral ventricle. Somata of a group of cells around the central canal (boxed area) are specifically labelled (magenta). Punctate signal below are ventral projections from these cells. Dotted line demarcates the boundary between gray and white matter. aii, Magnified view of the boxed area, showing the overlap between AAV labelling (magenta) and immunohistochemical signal for PKD2L1 (green), a known marker of CSF-cNs. Nuclear stain by DAPI is in blue. **b**, Morphology of labelled CSF-cNs (green). Cell bodies of the labelled cells (arrows) extend fine projections into the central canal (CC, dotted line). Arrowheads indicate the terminal enlargement of these projections, a stereotypic morphological feature of CSF-cNs. Ependymal cells are identified by their strong DAPI signal (blue). **c**, Projections from CSF-cNs to ventral white matter, as revealed by AAV-mediated PLAP expression and subsequent NBT/BCIP staining. **d**, Heat map showing the transcript expression level of known CSF-cN markers (*Pkd2l1, Gad1, Gad2*) and that of the κ opioid receptor, *Oprk1*. Color scale is based on median-of-ratios calculation by DESeq2. TdTomato-labelled CSF-cNs were dissociated and enriched by FACS from 4 different mice for bulk mRNA-sequencing. Each column represents one such sample. CSF-cN markers and Oprk1 share similar expression profiles as tdTomato transcript across samples, suggesting expression of these genes in CSF-cNs. The expression of the housekeeping gene *Actb* remains constant across samples. **e**, Immunohistochemical detection of OPRK1 (magenta) in CSF-cNs (green). Merged signal (white) on top; images from individual fluorescence channels at the bottom. **f**, Expression pattern of *Pdyn*, which encodes an OPRK1 ligand, in the spinal ependymal region of a *Pdyn*^*Cre*^*;Rosa26*^*LSL-tdTomato*^ mouse. *Pdyn*^*+*^ cells (magenta) are located close to CSF-cNs (green). **g**, Immunostaining confirmation of protein PDYN expression (green) in *Pdyn*^*Cre*^ labelled cells (magenta). Scale bars for **e-g** are 20 μm.

### κ opioid receptor and ligand are expressed in the ependymal region

Gaining specific genetic access to CSF-cNs enabled us to obtain clues about their potential function by transcriptome profiling. We expressed tdTomato in CSF-cNs of 4 wildtype C57BL6/J mice by AAV injection, then captured the tdTomato^+^ cells by fluorescence-activated cell sorting (FACS), generated independent cDNA libraries, and performed bulk sequencing. As reflected by the presence of glial transcripts (e.g., proteolipid protein 1 from oligodendrocytes and purinergic receptor p2y12 from microglia), our cell preparations were apparently contaminated to different degrees by glial fragments that likely adhered to the surface of tdTomato^+^ cells. Pertinently, *tdTomato* transcripts were also represented at different levels across the 4 samples (Fig. 1d, top row). Using this as a reference, we identified genes showing a relative abundance across samples matching that of *tdTomato*, thereby zeroing in on transcripts specifically enriched in CSF-cNs. Indeed, known CSF-cN markers, such as *Pkd2l1, Gad1* and *Gad2* (10, 20), were among the identified genes (Fig. 1d), together with those of relevance to neurotransmission and GABA metabolism (Supplementary Fig. 2a and b). Intriguingly, expression of the κ opioid receptor gene, *Oprk1*, correlated strongly with that of *tdTomato* transcript (Fig. 1d), suggesting that it is reasonably enriched in CSF-cNs. Consistent with this, OPRK1 protein was readily detected in CSF-cNs by immunohistochemical staining (Fig. 1e).

Next, we asked whether there is any κ opioid ligand in the vicinity of CSF-cNs. Prodynorphin (PDYN) is the precursor for several endogenous OPRK1 peptide ligands, including dynorphin A (DYNA), dynorphin B and big dynorphin (21). We examined spinal cord sections of *Pdyn*^*Cre*^*;Rosa26*^*LSL-tdTomato*^ mice injected i.c.v. with AAV2/2-GFP, in which *Pdyn*^*+*^ cells were labelled by tdTomato and CSF-cNs by GFP. Interestingly, in addition to previously reported *Pdyn*^*+*^ cells in the dorsal horn (22) (Supplementary Fig. 1h, left), we observed prominently labelled cells at the two poles of the ependyma, immediately apposing CSF-cNs (Fig. 1f). Immunostaining with an antibody against DYNA also highlighted the same region around the central canal (Fig. 1g), confirming expression at the protein level. We made use of the *MORF3* mouse line (23) to label the ependymal *Pdyn*^*+*^ cells sparsely for morphological tracing. In coronal spinal cord sections, *Pdyn*^*+*^ cells located at the dorsal pole of the ependyma extended a long process dorsally across the border between gray and white matters (Supplementary Fig. 1f and g), sometimes reaching all the way to the edge of the spinal cord. Ventral *Pdyn*^*+*^ cells projected ventrally instead, arborizing in the same field where CSF-cNs’ projections were found (Supplementary Fig. 1f and g). Both *Pdyn*^*+*^ cell types morphologically resembled radial glia but appeared to be negative for the astroglial marker GFAP. Instead, they were positive for the transcription factor SOX2 and the intermediate filament protein NESTIN (Supplementary Fig. 1h), two common markers of neural stem cells and progenitors previously shown to mark a subset of ependymocytes (24).

In all, we have identified a novel population of PDYN^+^ cells in the ependymal region that express κ opioids and reside near CSF-cNs that express the κ opioid receptor, hinting at some neurochemical communication between the two cell types.

### Activation of κ opioid receptor excites CSF-cNs

Activation of OPRK1 is typically believed to trigger an inhibitory cellular response through a Gα_i_ or Gβγ-mediated pathway (25, 26). However, there are few suggestions of excitatory responses such as in cultured astrocytes involving L-type voltage-gated calcium channels (Ca_v_ channels) (27, 28), as well as in the rat nucleus raphe magnus involving the mobilization of internal calcium stores (29). We were curious to see whether OPRK1 stimulation would inhibit or enhance the activity of CSF-cNs. We started by performing current-clamp recording on individual tdTomato-labelled CSF-cNs in acutely harvested spinal cord slices. At near resting membrane potential (−60 mV), CSF-cNs exhibited spontaneous action potentials at baseline as previously reported, owing to the spontaneous opening of PKD2L1 channels (11, 15). Interestingly, local application of the κ opioid ligand, DYNA_1-17_ (1 μM), increased the firing frequency above baseline (Fig. 2a, black). In addition to OPRK1, κ opioid ligands have been suggested to act on bradykinin receptors (30), NMDA receptors (31, 32) and/or delta and mu opioid receptors (33). Although none of these other candidate receptors are enriched in CSF-cNs based on our transcriptomic data (Supplementary Fig. 2c), we nevertheless tested whether OPRK1 is required for the response of CSF-cNs to κ opioid ligands. Indeed, the DYNA-evoked response was blocked when a selective OPRK1 antagonist, Nor-BNI (0.1 μM), was bath applied (Fig. 2a, red).

**Figure 2.**
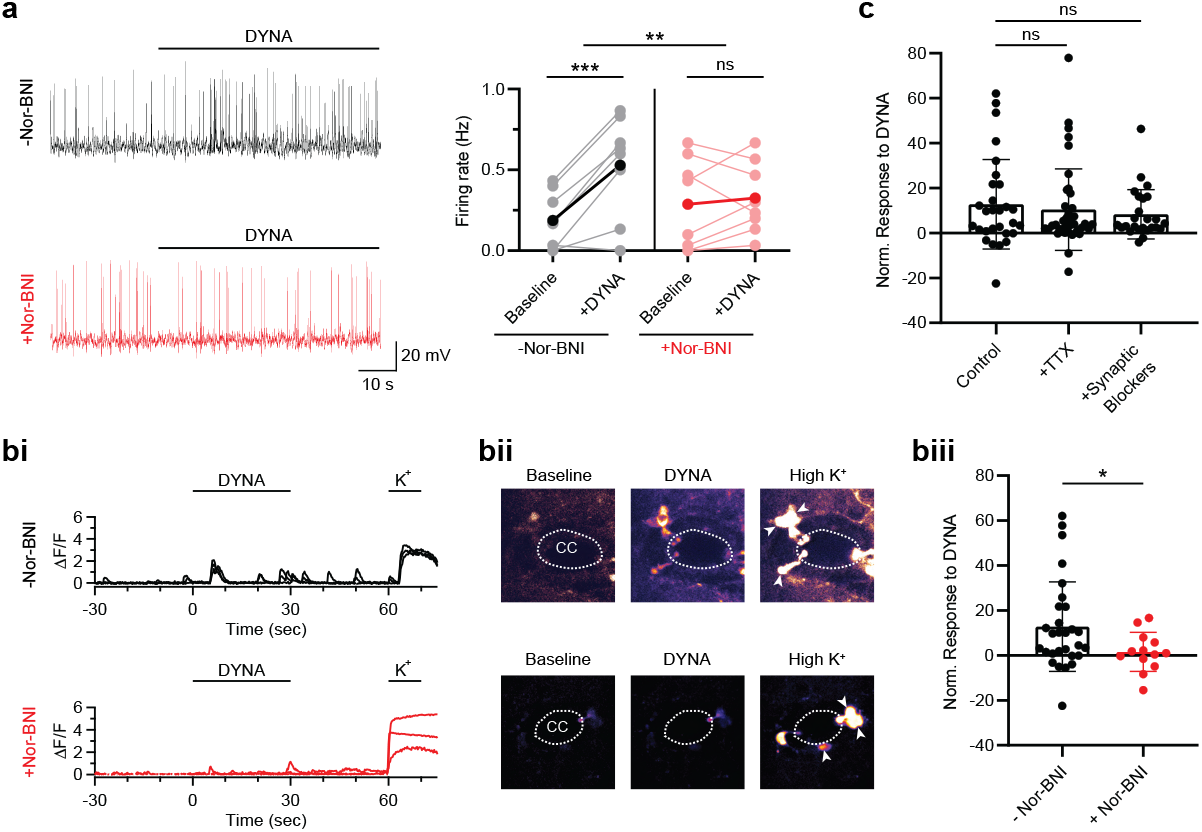
Activation of κ-opioid receptor excites CSF-cNs. **a**, OPRK1-dependent electrical responses of CSF-cNs to the κ opioid ligand, DYNA. Representative current-clamp recording traces showing spontaneous and DYNA-evoked firing in CSF-cNs in the absence (black) and presence (red) of an OPRK1 inhibitor, Nor-BNI. Collective data in bar graph on right. Individual cells are in light colors and averages in dark colors. Two-way ANOVA with Šidák correction for multiple comparisons, **p *≤* 0.01; ***p *≤* 0.001; ns: not significant, n = 8 and 8. **b**, OPRK1-dependent Ca^2+^ responses of CSF-cNs to DYNA. **bi**, Representative ΔF/F traces showing the responses of GCaMP5G-expressing CSF-cNs to local DYNA application in the absence (black) or presence (red) of Nor-BNI. Local application of a high K^+^ solution was used to reveal all responsive neurons in spinal cord slices. Each trace is from a single cell. **bii**, ΔF/F images for the cells (arrowheads) in bi. Images are temporal averages over 10 sec of baseline or for the duration of the stimuli. CC: central canal. **biii**, Collective data comparing responses of CSF-cNs to DYNA in the absence (black) and presence (red) of Nor-BNI. Each dot shows the integral DYNA response of a single cell normalized to the high-K^+^ response (see Methods). Welch’s t test, *p *≤* 0.05, n = 29 and 13. **c**, DYNA-evoked Ca^2+^ responses under bath application of TTX or synaptic blockers. Post-hoc Dunnett tests for comparison with control, which is same as the – Nor-BNI condition in biii, ns: not significant, n = 29, 39 and 25.

We confirmed the excitatory effect of OPRK1 activation by performing calcium imaging of GCaMP5G-expressing CSF-cNs in spinal cord slices from *Tg(PKD2L1-Cre);Polr2a*^*GCaMP5G-tdTomato*^ mice. In response to DYNA (1 μM), a considerable number of CSF-cNs increased their calcium activity significantly above baseline (Fig. 2b). Two other OPRK1 agonists, Nalfurafine (1 μM) and BRL-52537 (10 μM), also elicited similar calcium responses in many CSF-cNs (Supplementary Fig. 3). We noticed that not all CSF-cNs responded, and those that did respond continued to spike even after stimulus withdrawal, likely reflecting inefficient perfusion of agents into and out of the tissue slices. However, it was clear that the responses were OPRK1-dependent since they were blocked by the OPRK1 antagonists, Nor-BNI (0.1 μM, Fig. 2b and Supplementary Fig. 3b) or DIPPA (1 μM, Supplementary Fig. 3a).

To ascertain whether the response to DYNA is intrinsic to CSF-cNs or synaptically driven, we included the broad-spectrum voltage-gated sodium (Na_v_) channel blocker, tetrodotoxin (TTX), in the bath to block action potential firing in upstream neurons. The response of CSF-cNs to DYNA persisted in the presence of TTX (Fig. 2c). Previously, CSF-cNs were found to express functional AMPA/kainate-type ionotropic glutamate receptors (15), GABA_A_ receptors (11), nicotinic acetylcholine receptors (15, 34) and glycine receptors (11). We therefore also applied a cocktail of synaptic blockers to inhibit fast glutamatergic, GABAergic, cholinergic and glycinergic neurotransmission. Again, no significant effect was seen on DYNA-evoked responses (Fig. 2c). Moreover, DYNA did not influence the rate of spontaneous postsynaptic events in voltage-clamp recordings of CSF-cNs (Supplementary Fig. 5e). Together, our experiments suggest that a functional excitatory OPRK1 signaling pathway is intrinsic to CSF-cNs.

We next used pharmacological tools to gain initial insights into how OPRK1 activation regulates CSF-cN excitability. A canonical GPCR-mediated transduction pathway that promotes a rise in intracellular calcium involves activation of G_q_-phospholipase C (PLC) signaling. Indeed, a Gα_q_ inhibitor (YM254890, 10 μM) diminished DYNA-evoked calcium responses in CSF-cNs (Supplementary Fig. 4a, blue region). Inhibition was also observed with the PLC inhibitor, U73122 (10 μM) but not with its inactive analog (U73343, 10 μM) (Supplementary Fig. 4a, blue region). PLC catalyzes the hydrolysis of phosphatidylinositol 4,5-bisphosphate (PIP_2_) to inositol 1,4,5-triphosphate (IP_3_) and diacylglycerol (DAG). IP_3_ can elicit a release of calcium from intracellular stores, which subsequently opens calcium-sensitive membrane ion channels. In fact, in rat nucleus raphe neurons, regulation of HCN channels by internal calcium has been proposed to explain the excitatory response triggered by OPRK1 stimulation (29). In CSF-cNs, however, we observed no significant effect of the HCN blocker, Ivabradine (10 μM), on DYNA responses (Supplementary Fig. 4a, yellow region). Moreover, depletion of internal calcium stores with thapsigargin (4 μM) (Supplementary Fig. 4a, yellow region) had a statistically significant, but only minor effect. An alternative pathway downstream of PLC involves the activation of TRPC channels by DAG. CSF-cNs express *Trpc1* and *Trpc6* transcripts (Supplementary Fig. 2d) and are particularly abundant in PKD2L1, a TRPP channel that may also be positively regulated by PLC (35). Nevertheless, we did not detect any macroscopic current or an increase in single-channel opening events upon DYNA application by voltage-clamp recording of CSF-cNs (at −80 mV, Supplementary Fig. 5a-d), arguing against membrane depolarization through these non-selective cation channels.

Finally, we turned to the possibility of DAG activating protein kinase C (PKC), which is known to modulate the activity of Ca_v_ channels via phosphorylation, consequently shifting their voltage dependence and/or kinetic properties (36). Indeed, the cell-permeable PKC inhibitor, chelerythrine chloride (10 μM), reduced DYNA-evoked calcium responses in CSF-cNs (Supplementary Fig. 4a, green region). Removal of extracellular calcium also weakened the DYNA response (Supplementary Fig. 4a, green region). Further experiments involving individual Ca_v_ channel blockers suggest that external calcium enters CSF-cNs through P/Q-, N- and T-type Ca_v_ channels (i.e., Ca_v_2.1, 2.2 and 3, respectively) and less so through L-type (Ca_v_1.x) channels (Supplementary Fig. 4a, green region). Consistent with this, our bulk sequencing data indicates that certain Ca_v_2 and 3 channel subunits are enriched in CSF-cNs (Supplementary Fig. 2d). Previous pharmacological studies have also suggested roles for T-type and HVA calcium channels (which include P/Q- and N-type) in mediating spontaneous and evoked spiking in CSF-cNs (16). Of note, our earlier recordings did not reveal the engagement of Ca_v_ channels because the holding potential (−80 mV) was too negative for channel activation, even after a shift of voltage sensitivity by phosphorylation. However, as noted above, CSF-cNs are known to have a resting membrane potential of ∼ −55 mV and exhibit PKD2L1-mediated spontaneous action potentials (11, 15), which may provide the depolarization needed to drive Ca_v_ channel activation.

In sum, our pharmacological data suggest that OPRK1 activation of CSF-cNs mainly involves a G_q_-PLC-PKC cascade, leading ultimately to increased calcium influx through Ca_v_ channels and action potential firing (Supplementary Fig. 4b).

### Tonic activation of CSF-cNs by κ opioids suppresses ependymal proliferation

What action does κ signaling exert in the ependymal region? As noted above, ependymal proliferation can be inhibited by GABA (8). CSF-cNs are well poised to be the cellular origin of this GABA-mediated regulatory mechanism because they are GABAergic (20, 37, 38) and reside near ependymal cells. Hence, we tested whether ependymal cell proliferation would be affected by interfering with κ signaling of CSF-cNs *in vivo*. We injected healthy adult wildtype C57BL6/J mice with a κ agonist (Nalfurafine, 20 μg/kg, i.p.) or antagonist (Nor-BNI, 10 mg/kg, i.p.) daily for 8 consecutive days. Control mice were injected with the respective vehicles. We then subjected the mice to an overlapping but staggered week of treatment with EdU (50 mg/kg, i.p.), a thymidine analog that is incorporated into newly synthesized DNA during cell division (Fig. 3a). After that, we prepared spinal cord sections from these animals and counted the number of EdU^+^ ependymal cells. We found that the average EdU^+^ cell count was not significantly affected by the κ agonist. However, a slight decreasing trend was seen when cell counts from each section were tallied as a distribution histogram (Fig. 3bii, left). Most likely, the low number of proliferating ependymal cells at baseline makes it difficult to detect small reductions. On the other hand, the κ antagonist significantly upregulated the number of EdU^+^ cells in the spinal ependymal region, as evident from both representative histological images (Fig. 3bi, right) and collective data (Fig. 3bii, right). Thus, we conclude that a constitutively active κ signaling pathway normally suppresses ependymal cell proliferation.

**Figure 3.**
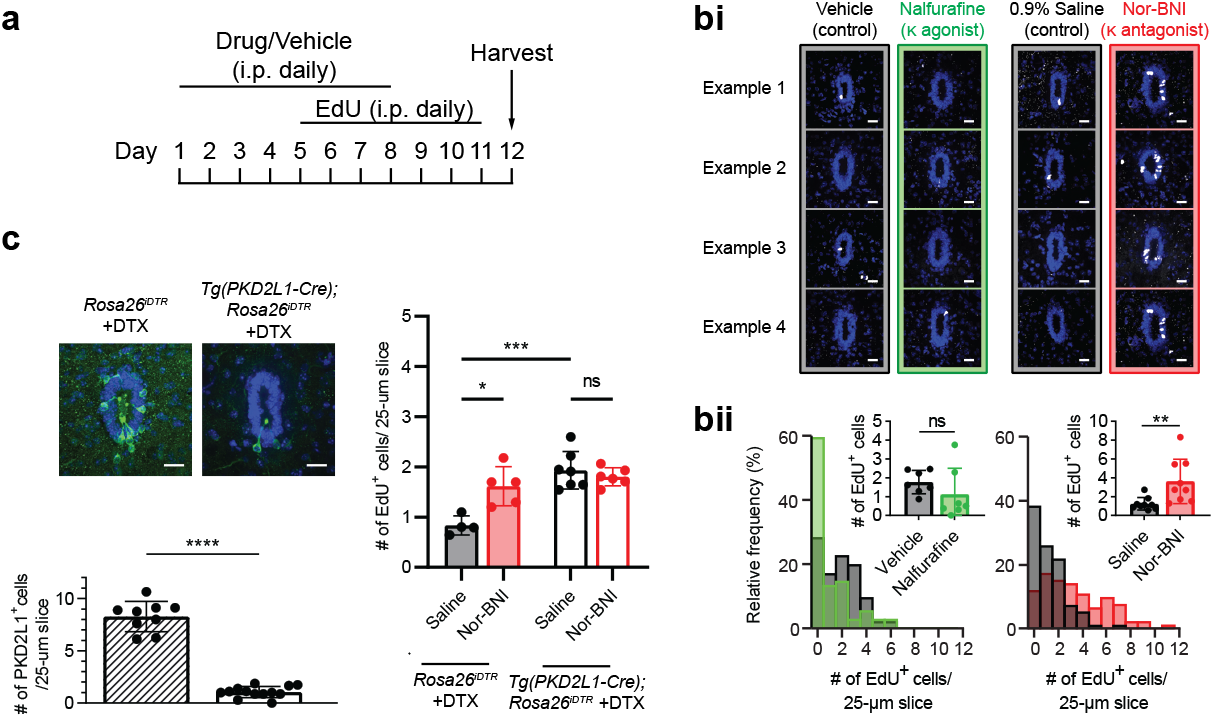
Constitutive κ signaling via CSF-cNs suppresses ependymal cell proliferation *in vivo*. **a**, Drug injection scheme for examining changes in the number of EdU-labelled proliferating cells in spinal ependymal region after systemic treatment with κ agonist (20 μg/kg Nalfurafine), antagonist (10 mg/kg Nor-BNI) or respective vehicles. **bi**, Representative images of EdU^+^ cells (white) in the ependyma (blue) in 4 different spinal cord sections from a mouse from each treatment group. **bii**, Distribution histogram showing the frequency of spinal cord sections (25-μm) containing the indicated number of EdU^+^ cells. (Inset) Bar graph summary of the same data. Each dot represents the average of multiple sections from a single mouse. Welch’s t test, **p *≤* 0.01; ns: not significant, n = 7 (vehicle), 7 (Nalfurafine), 9 (saline) and 9 (Nor-BNI). **c**, Dependence of Nor-BNI’s effect on CSF-cNs. (Left) Representative images of spinal cord ependymal region from DTX-injected *Tg(PKD2L1-Cre);Rosa26*^*iDTR*^ and control mice immunostained with PKD2L1 antibody to reveal CSF-cNs. Scale bars are 20 μm. Collective data below show near complete ablation of CSF-cNs in experimental animals. Each dot is the average cell count from multiple spinal cord slices from a single mouse. Welch’s t test, ****p *≤* 0.0001, n = 9 and 13. (Right) Number of EdU^+^ cells in saline versus Nor-BNI-treated mice with and without CSF-cN ablation. Data quantified as in bii. One-way Welch ANOVA with Dunnett’s correction for multiple comparisons, *p *≤* 0.05; ns: not significant, n = 4, 5, 7, 6.

To determine whether the robust antagonist effect depends on CSF-cNs, we ablated these neurons by generating *Tg(PKD2L1-Cre);Rosa26*^*iDTR*^ mice in which the diphtheria toxin receptor (DTR) is selectively expressed in CSF-cNs. Diphtheria toxin (DTX, 50 μg/kg, i.p.) was administered for 5 consecutive days prior to the same antagonist and EdU injection scheme described above. Immunostaining with a PKD2L1 antibody verified that most CSF-cNs in these animals were eliminated compared to DTX-treated control *Rosa26*^*iDTR*^ mice that did not carry the Cre allele (Fig. 3c, left). As in wildtype mice, the κ antagonist Nor-BNI enhanced ependymal proliferation in control *Rosa26*^*iDTR*^ animals, although to a lesser extent possibly due to some DTX side effects (Fig. 3c, right bar graph, bars 1 and 2). When CSF-cNs were ablated, vehicle-injected animals showed significantly more EdU^+^ ependymal cells (Fig. 3c, right bar graph, bars 1 and 3), demonstrating that CSF-cNs exert constant suppression on ependymal proliferation under normal physiological conditions. Moreover, the proliferation-enhancing effect of the κ antagonist was lost after CSF-cN ablation (Fig. 3c, right bar graph, bars 3 and 4), further supporting the notion that CSF-cNs are necessary for κ opioid-mediated suppression of proliferation. This overall suppressive effect is also consistent with our finding described above that κ opioids exert an excitatory action on CSF-cNs, which are known to be inhibitory (i.e., GABAergic) (20, 37, 38).

### κ agonist reduces injury-induced ependymal proliferation

Injury to the midline of the spinal cord has been shown to elicit extensive ependymal cell proliferation, which is critical for scar formation (3–7). Glial scars are a double-edged sword: they help to prevent further injury to the spinal cord but also present a barrier to axonal regeneration (1). We asked whether systemic administration of κ opiate drugs can be used as a means for controlling the degree of ependymal cell proliferation, which may provide therapeutic benefits. We examined this in a common injury model where the dorsal aspect of the spinal cord is transected (i.e., dorsal hemisection) (Fig. 4a). To best capture the early phase of injury-induced proliferation, we adopted a more compact injection scheme by moving the beginning of the EdU doses forward to the day after injury (Fig. 4b), which was also the second day of drug treatment. We confirmed that sham operated controls behaved similarly to uninjured wildtype mice in that the κ agonist Nalfurafine has no impact on ependymal proliferation (Fig. 4c). Consistent with earlier reports (3–6), dorsal hemisection caused a substantial increase in the number of EdU^+^ ependymal cells in mice injected with vehicle relative to sham controls (Fig. 4c). Remarkably, this injury-induced proliferation could be lowered by ∼ 55% with Nalfurafine treatment (Fig. 4c), indicating that κ agonists indeed have a pronounced suppressive effect on ependymal cell proliferation.

**Figure 4.**
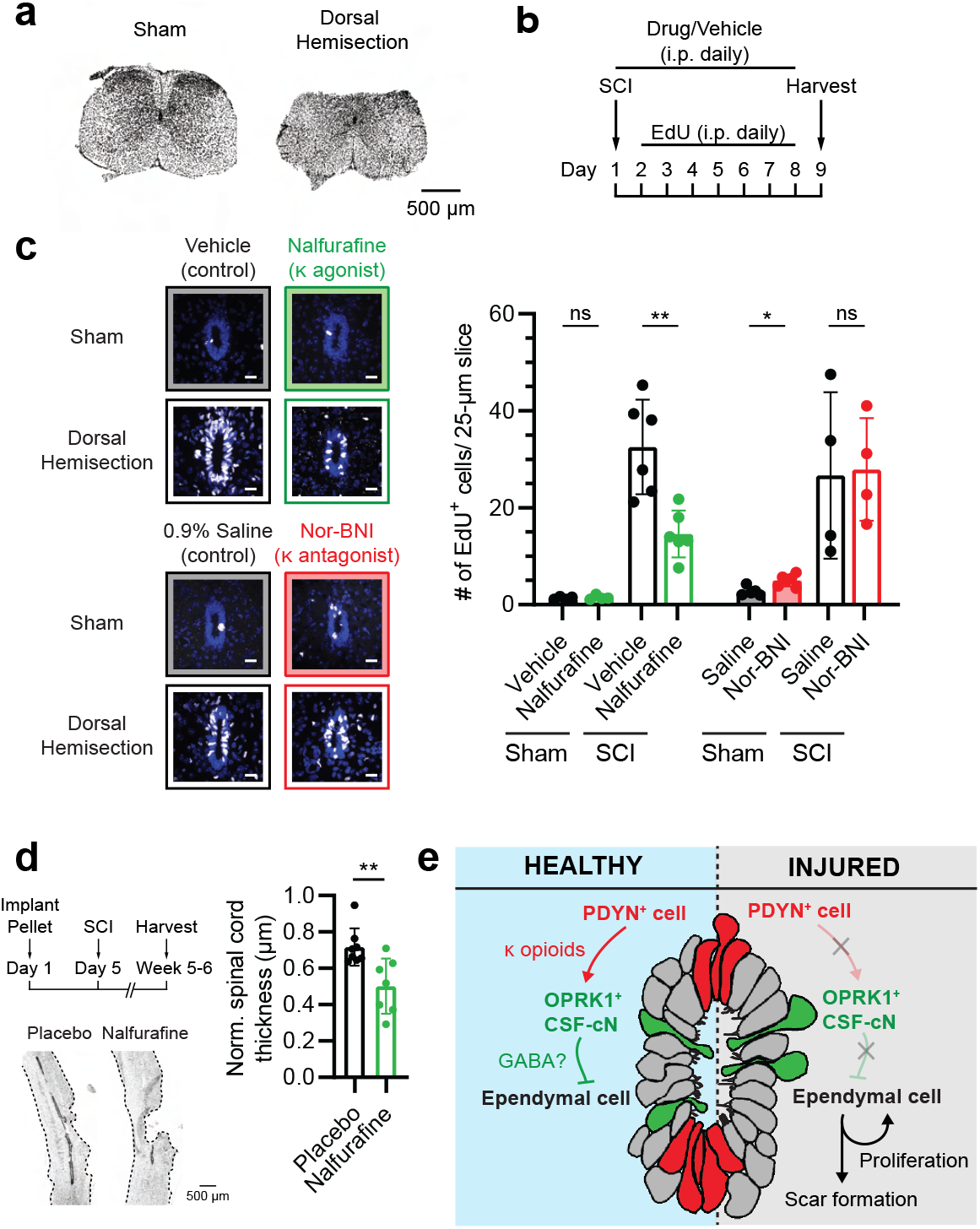
Systemic administration of κ agonist reduces ependymal proliferation induced by spinal cord injury. **a**, DAPI-stained coronal sections of spinal cords from a sham-operated mouse and a mouse with dorsal hemisection to demonstrate the degree of injury. **b**, Injury and drug treatment regimen. **c**, Number of EdU^+^ cells in sham-operated animals and mice subjected to dorsal hemisection with or without systemic treatment with κ agonist (20 μg/kg Nalfurafine) or antagonist (10 mg/kg Nor-BNI). Representative images on left and bar graph summary on right. Welch’s t test, *p *≤* 0.05; **p *≤* 0.01; ns: not significant, n = 4, 4, 6, 6, 5, 6, 4, 4. Scale bars are 20 μm. **d**, Effect of long-term κ agonist administration (Nalfurafine, 0.027mg/pellet with 60-day release rate, 5-6 weeks of treatment) on thickness of hemisected spinal cords (outlined in dashes). Spinal cord thickness was minimum measurements taken from injured regions normalized to that from uninjured regions in the same sagittal sections. Welch’s t-test, **p *≤* 0.01, n = 8 and 7. **e**, Schematic summary of the proposed signaling pathway. In healthy conditions, PDYN^+^ cells release κ opioids to activate OPRK1^+^ CSF-cNs, which in turn suppress ependymal proliferation. Injury to PDYN^+^ cells prevents opioid peptide release and subsequent suppression by CSF-cNs, thereby allowing ependymal cells to proliferate.

In a complementary set of experiments, we examined the effect of the κ antagonist Nor-BNI. In sham operated controls, we again saw a significant increase in the number of EdU^+^ ependymal cells in antagonist-versus vehicle-treated mice (Fig. 4c); this trend was similar to, but perhaps slightly weaker than that seen in the healthy condition, likely reflecting the shorter drug treatment (1 versus 4 days) preceding EdU administration in the modified experimental paradigm. Importantly, after dorsal hemisection, ependymal cells proliferated dramatically and Nor-BNI did not further enhance this proliferation, suggesting that the constitutive κ signaling that occurred under healthy condition had now been interrupted by injury. We note that injury-induced proliferation was much greater than that induced by κ antagonist in healthy mice. This may partly reflect limited antagonist dosage or involvement of other molecular signals in the injury response. Incidentally, CSF-cNs and PDYN^+^ cells remain non-proliferative after injury (Supplementary Fig. 6a and b).

Genetic suppression of ependymal proliferation has been shown to cause secondary enlargement of the lesion site in injured spinal cords (5). We asked whether pharmacological suppression of ependymal proliferation by long-term application of a κ agonist would produce a similar effect. For this, we implanted mice subcutaneously with either a placebo or Nalfurafine pellet (0.027 mg Nalfurafine per pellet, 60-day release rate) and measured the spinal cord thickness 5-6 weeks after dorsal hemisection. Indeed, the injured segment of the spinal cords from Nalfurafine-treated animals showed less buildup of GFAP^+^ scar (Supplementary Fig. 6c) and very severe thinning (Fig. 4d) compared to those from controls, consistent with the notion that opioid regulation of ependymal proliferation is critical for supporting tissue integrity after injury.

## Conclusions

Our findings provide insight into how the proliferation of spinal cord ependymal cells is regulated *in vivo* (Fig. 4e). In normal healthy conditions, CSF-cNs receive constant κ opioid stimulation, likely from neighboring PDYN^+^ cells. Activation of the κ opioid receptor in CSF-cNs excites these neurons, which release GABA and/or other inhibitory signals that suppress ependymal cell proliferation. Upon spinal cord injury, constitutive κ signaling halts, possibly due to physical damage to the radial processes of PDYN^+^ cells. As a result, suppression by CSF-cNs is relieved and ependymal cells are allowed to proliferate to facilitate scar formation. The implication of radial glia-like PDYN^+^ cells in this process may explain why ependymal proliferation is triggered most prominently when the injury occurs close to the midline, rather than laterally or by a general crush (4, 39). The neurotransmitter GABA has been shown to elicit GABA_A_ receptor-mediated electrical responses in ependymal cells (40) and restrict their proliferation (8), thus representing a likely suppressive signal released by GABAergic CSF-cNs. Interestingly, CSF-cNs express functional GABA_A_ receptors themselves (11) and hence they may possess a feedback mechanism for autoregulating GABA release. CSF-cNs also express several other neurotransmitter receptors (11, 15, 34), although their upstream synaptic partners are largely uncharacterized. It remains to be determined whether ependymal proliferation is also modulated by synaptic input to CSF-cNs under specific physiological circumstances. Moreover, CSF-cNs have been reported to be part of some motor circuits (13, 17, 19) and it will therefore be interesting to see whether these neurons coordinate locomotion readjustment after spinal cord injury.

κ agonists are known to reduce pain and itch behaviors by curbing neurotransmission from primary sensory afferents to spinal cord dorsal horn neurons and inhibiting peripheral neurogenic inflammation (41). In fact, the agonist Nalfurafine used in the current study has already been approved in several countries as an anti-pruritic (42). Our work now raises the important question of whether κ opioid receptor drugs can also be employed to control ependymal cell proliferation, thus modulating scar formation and promoting wound healing in certain types of spinal cord injuries. It is well accepted that glial scars present a non-permissive physical and chemical environment for axon regrowth. However, accumulating evidence suggests that scar formation also serves a key protective function (1, 2). In a previous study (5), ablation of ependymal cell proliferation through genetic knockout of cell cycle genes rendered spinal cords more susceptible to secondary tissue damage and neuronal loss following injury. We also observed a similar thinning of spinal cord tissues after long-term κ agonist treatment to suppress ependymal proliferation post injury. Hence, a dynamic control of the degree of scar formation may have therapeutic utility. Future studies will determine whether κ opiate drugs can be harnessed to achieve such regulation in addition to providing pain relief.

## Materials and Methods

### Animals

All animal experiments were conducted in accordance with protocol AN192533 approved by the Institutional Animal Care and Use Committee, University of California – San Francisco. Mice were raised under regular diurnal (12:12) light-dark cycles with ad libitum access to food and water. We used mice of both sexes between the age of 5-12 weeks. *Tg(PKD2L1-Cre)* mice (MGI: 5691401)(10) were kindly provided by Dr. Charles Zuker (Columbia University). Wildype C57BL6/J mice as well as *Rosa26*^*LSL-tdTomato*^ (MGI: 3813512) (43), *Polr2a*^*GCaMP5G-tdTomato*^ (MGI: 5560331) (44), *MORF3* (MGI: 292704) (23) and *ROSA26*^*iDTR*^ (MGI: 3772576) (45) mice were acquired from Jackson Laboratory. For cell ablation, 1 mg/ml stock solution of DTX (List Biological Laboratories, 150) was prepared with water, aliquoted and stored at −20°C until use. When needed, the DTX solution was diluted to 5 ng/μl with sterile 0.9% saline and 50 μg/kg was administered i.p. daily for 5 consecutive days. Littermates were used as genotype controls whenever possible; when not, age-matched animals were used.

### AAV packaging and intracerebroventricular injection

AAV2 or PHP.eB viral preparations made from pAAV-CAG-tdTomato (59462) or pAAV-CAG-GFP (37825) plasmids were purchased from Addgene. AAV2 was generated from pAAV-FLEX-PLAP plasmid (Addgene, 80422) by triple transfection into HEK293FT cells, followed by purification of AAV particles with the iodixanol gradient ultracentrifugation method. Viral titer was standardized to ∼2E12 GC/ml for *in vivo* delivery.

Under isoflurane anesthesia, a mouse was positioned on a heating pad on a stereotaxic instrument (David Kopf Instruments, Model 1900). Hairs were removed from the top of the head and the surgical site was disinfected with alternating scrubs of 7.5% Povidone Iodine (Betadine surgical scrub) and 70% Ethanol. Lidocaine (Vedco, NDC 50989-417-12, 1.6 mg/ml in sterile 0.9% saline, 100 μl/mouse) was injected subcutaneously prior to making a ∼1-cm long mid-sagittal skin incision over the skull. The bregma and lambda points were identified, and the skull was levelled by using a stereotaxic alignment indicator (Kopf Instruments, Model 1905) according to the manufacturer’s instructions. A small hole was opened on the skull at the coordinates (X = +1.7, Y = −0.9) or (X = +1.0, Y = −0.4) by using a stereotaxic drill (Kopf Instruments, Model 1911) coupled to a 0.027” ball drill bit (CircuitMedic, 115-6050). The cranial window was irrigated with sterile PBS immediately after opening to prevent dehydration. A Wiretrol II glass pipette (Drummond Scientific Company, 5-0002010), pulled to give a sharp tip (Sutter, P-97 micropipette puller) before the experiment, was mounted onto the stereotaxic frame and was loaded with 5 μl of AAV solution. The pipette tip was slowly advanced to 2.2 mm beneath the brain surface and the AAV solution was dispensed gradually by manually controlling the Wiretrol plunger. Tissues were allowed to take up the AAV solution for 10 min before the pipette was retrieved slowly. Finally, the skin was closed by suturing and the mouse was allowed to recover on a warm pad. Buprenorphine (Par Pharmaceutical, NDC 42023-179-05, 0.1 mg/kg) and Meloxicam (Pivetal, NDC 46066-937-13, 10 mg/kg) were administered i.p. during and after the surgery for analgesia.

### Immunohistochemistry and Click-iT EdU imaging

Mice were euthanized by CO_2_ asphyxiation according to American Veterinary Medical Association’s guidelines. Transcardiac perfusion with phosphate-buffered saline (PBS, Quality Biological, 119-069-491) followed by 4% paraformaldehyde (PFA, Electron Microscopy Sciences, 15714) was performed immediately after. Spinal cords were harvested into 4% PFA and fixed overnight. After 3 washes with PBS, spinal cords were transferred to 30% sucrose (Sigma-Aldrich, S7903) and allowed to settle at 4°C. The tissues were then cryopreserved in Tissue-Tek O.C.T. Compound (Sakura Finetek USA) and sectioned at a thickness of 25-50 μm on a Leica CM3050 S cryostat. Spinal cord sections were collected in PBS and were stored in freezing medium (30% w/v sucrose [Sigma-Aldrich, S7903], 1% w/v polyvinylpyrrolidone [Sigma-Aldrich, PVP40], 30% v/v ethylene glycol [Sigma-Aldrich, 102466] in 50 mM phosphate buffer) at −20°C until use. On the day of experiment, sections were retrieved into PBS and were washed with PBST (i.e., PBS containing 0.5% Triton X-100 [Sigma-Aldrich, T8787]) for multiple times. Subsequently, sections were incubated with blocking buffer (PBST containing 10% normal goat serum [Thermo Fisher Scientific, 16-210-064] or donkey serum [Sigma-Aldrich, D9663]) at room temperature for 1 hr. Overnight primary antibody incubation was done at 4°C with one or more of the following antibodies in blocking buffer: rabbit anti-DsRed (Takara Bio, 632496, 1:500), goat anti-mCherry (Sicgen, AB0081-200, 1:500), rabbit anti-PKD2L1 (Sigma-Aldrich, AB9084, 1:500-700), rabbit anti-OPRK1 (Abcam, ab183825, 1:500), chicken anti-GFP (Abcam, ab13970, 1:500), rabbit anti-DYNA (Peninsula Laboratories, IHC8676, 1:500) and chicken anti-GFAP (Abcam, ab4674, 1:500). On the following day, sections were washed with PBST for 3 times and incubated with the appropriate secondary antibodies at 1:500 in blocking buffer. Secondary antibodies include goat anti-rabbit IgG-Alexa Fluor 568 (Thermo Fisher Scientific, A-11036), donkey anti-goat IgG-Alexa Fluor 568 (Thermo Fisher Scientific, A-11057), goat anti-rabbit IgG-Alexa Fluor 488 (Thermo Fisher Scientific, A-11034) and goat anti-chicken IgY-Alexa Fluor 488 (Thermo Fisher Scientific, A-11039). The nuclei stain, 4,6-diamidino-2-phenylindole (DAPI, 0.5 μg/ml, Thermo Fisher Scientific, D1306), was included in one of the final PBST washes. Lastly, sections were laid on a glass slide and mounted with Fluoromount-G (SouthernBiotech, 0100-01). Z-stack images were taken with a Nikon CSU-W1 spinning disk confocal microscope or a Nikon Ti Inverted Microscope (UCSF Center for Advanced Light Microscopy). Image stitching was done by NIS-Elements software (Nikon). Maximal intensity projections were generated by the Fiji software (NIH). For quantifying GFAP immunosignal intensity, a segmented line was drawn along the injured site and pixel intensity was measured by the Fiji software.

To detect EdU^+^ cells, 25-μm cryosections of the spinal cord were made as above and processed through the aforementioned immunostaining procedure when necessary. After the final PBST wash, the freely-floating sections were incubated in the Click-iT reaction cocktail (Thermo Fisher Scientific, C10340) in the dark at room temperature with shaking for 1 hr. Following one PBST wash, the sections were incubated with DAPI, then further washed, mounted and imaged as described above.

### PLAP staining and tissue clearing

For PLAP staining of tissue sections, we prepared 50-μm cryosections of the spinal cord and brainstem from perfused mice as described above. To inactivate endogenous alkaline phosphatase activity, the samples were heated up in PBS in a 72°C water bath for 1 hr. The sections were washed twice in Buffer 1 (100 mM Tris base, 150 mM NaCl, pH 7.5) and twice in Buffer 2 (100 mM Tris base, 100 mM NaCl, 50 mM MgCl_2_, pH 9.5) before being transferred to the staining buffer (2% NBT-BCIP stock solution [Sigma-Aldrich, 11681451001] in Buffer 2). Samples were allowed to be stained for 30 min in the dark and were then dehydrated through a series of increasing methanol concentration (20%, 40%, 60%, 80%, 100%, room temperature, 30 min each). After mounting, images were taken with a Nikon Ti Inverted Microscope (UCSF Center for Advanced Light Microscopy). Stitched images were generated by NIS-Elements (Nikon Instruments).

For wholemount PLAP staining, spinal cords were harvested from perfused mice and fixed overnight in 4% PFA as described above. They were then bisected along the midline to expose the ependymal region. Heat inactivation and PLAP staining were carried out as above except that the incubation in staining buffer was 1 hr long. After dehydration, the tissues were stored in 100% methanol at room temperature until being transferred to 66% dichloromethane (DCM)/33% methanol on the following day for clearing at room temperature for 3 hrs on a rotator. Subsequently, methanol was washed away by two 15-min incubations in 100% DCM and the samples were allowed to stand and clear in a glass bottle filled completely with dibenzyl ether (DBE). The tissues were submerged in DBE in a home-made chamber when being imaged on an Olympus FV3000 microscope.

### Bulk mRNA-sequencing

Mice received intracerebroventricular injection of AAV2-CAG-tdT as described above. After 3-4 weeks, a mouse was perfused with ice-cold NMDG-aCSF (92 mM NMDG, 2.5 mM KCl, 10 mM MgSO_4_, 0.5 mM CaCl_2_, 1.2 mM NaH_2_PO_4_, 30 mM NaHCO_3_, 20 mM HEPES, 5 mM Na-ascorbate, 2 mM Thiourea, 3 mM Na-pyruvate, 25 mM Glucose, pH 7.4) and its spinal cord was freshly removed into the same solution by applying hydraulic pressure through a syringe to the open spine. The spinal cord was bisected along the midline with a stab knife (Surgical Specialties, 72-2201) on a sylgard (Dow, DC4019862) plate and embedded in 2% low-melting agarose. 200-μm thick sagittal slices containing the ependymal region were collected from each half of the spinal cord by using a Leica VT1000 S vibratome. Slices were cut into smaller pieces and transferred into Enzyme Mix 1 (955 μl of Buffer Z, 25 μl of Enzyme P [Miltenyi Biotec, 130-094-802]) in a 1.5-ml tube. After 15-min incubation in a 37°C water bath, Enzyme Mix 2 (15 μl of Buffer Y, 7.5 μl of Enzyme A [Miltenyi Biotec, 130-094-802]) was added for another 5-min, 37°C incubation. The digestion solution was then replaced with 1 ml HABG (97.75% Hibernate A Low Fluorescence [BrainBits, HALF], 2% B27 supplement [Thermo Fisher Scientific, 17504044], 0.25% GlutaMAX [Thermo Fisher Scientific, 35050061]). The digested tissues were triturated multiple times to mechanically dissociate the cells by using fire-polished glass pipettes with decreasing tip diameters. The cell suspension was then laid onto 4 ml of 20% Percoll (in HABG) and was centrifuged at 300 × rcf for 15 min in a cold room. The resulting supernatant was removed carefully, and the cell pellet was resuspended in 0.8 ml HABG containing Ribolock RNase inhibitor (Thermo Fisher Scientific, EO0382, 0.4 U/μl). A far-red LIVE/DEAD dye (Thermo Fisher Scientific, L34973, 1:1000) was added to the suspension for excluding dead cells during FACS. The cell suspension was sorted based on forward and side scatters, as well as tdTomato and LIVE/DEAD fluorescence in a BD Biosciences FACSAria II Cell Sorter (UCSF Laboratory for Cell Analysis). Fifty cells were collected in a tube containing ice-cold lysis buffer and processed for cDNA synthesis according to the Smart-Seq v4 manual (Takara Bio, 634890) with a PCR cycle number of 11. Subsequently, cDNA library was prepared by using Nextera XT DNA Library Preparation Kit (Illumina, FC-131-1024). We followed the manufacturer’s manual for tagmentation, amplification and cleanup steps, but estimated the concentrations of our libraries by comparing their SYBR green (Thermo Fisher Scientific, S7563) fluorescence to a standard curve made from λ DNA (Thermo Fisher Scientific, 25-250-010). Library quality was confirmed by using the Bioanalyzer High Sensitivity DNA Kit (Agilent Technologies, 5067-4626). Four libraries were pooled for pair-end sequencing with 100 cycles on a S1 flow cell lane in an Illumina NovaSeq 6000 System (UCSF Center for Advanced Technology). A technical replicate was included in a separate lane. About 40-60 million reads were obtained per sample and the sequencing quality was examined by using FASTQC. Adapters and low-quality reads were trimmed or removed by TrimGalore. Reads were then aligned to the annotated mouse reference genome (mm10) containing an additional exogenous sequence for AAV-CAG-tdT by using RSEM. Gene counts were imported into RStudio (v4.2.0) for analysis by using the DESeq2 module (v1.36.0, Bioconductor v3.16). Replicates were collapsed and the DESeq function was applied to normalize read counts. Results were displayed as heatmaps.

### Calcium imaging of spinal cord slices

A mouse was euthanized by CO_2_ asphyxiation, and its spinal cord was acutely harvested into ice-cold NMDG-aCSF (92 mM NMDG, 2.5 mM KCl, 10 mM MgSO_4_, 0.5 mM CaCl_2_, 1.2 mM NaH_2_PO_4_, 30 mM NaHCO_3_, 20 mM HEPES, 5 mM Na-ascorbate, 2 mM Thiourea, 3 mM Na-pyruvate, 25 mM Glucose, pH 7.4). After removing the meninges, the spinal cord was embedded in 2% low-melting agarose, and the thoracic-lumbar region was sectioned coronally at a thickness of 300 μm in ice-cold NMDG-aCSF by using a Leica VT1000 S vibratome. Each spinal cord slice was adhered to a small piece of filter membrane (Whatman Nuclepore Track-Etched Membrane, Sigma-Aldrich, WHA110657) for convenient handling. Slices were allowed to recover at 34°C in NMDG-aCSF for 30 min and then transferred to aCSF (124 mM NaCl, 2.5 mM KCl, 2 mM MgSO_4_, 2 mM CaCl_2_, 1.2 mM NaH_2_PO_4_, 24 mM NaHCO_3_, 5 mM HEPES, 12.5 mM Glucose, pH 7.4) for storage at room temperature until being used within 3 hrs. All solutions were constantly bubbled with 95% O_2_/5% CO_2_ to provide oxygen and maintain pH. When needed, a piece of filter membrane carrying a spinal cord slice was immobilized by vacuum grease in an imaging chamber and was placed under a Leica SP8 confocal microscope. The ependymal region was located under brightfield and CSF-cNs were identified by their tdTomato signal. Image acquisition was done by using the LAS X software (Leica Microsystems) at 4 Hz with a confocal pinhole equivalent to a depth of ∼ 220 μm of tissues. The tissues were bath perfused with bubbled room-temperature aCSF at a rate of ∼ 1 ml min^-1^. Pharmacological reagents were included in the bath and/or applied locally at a rate of ∼ 140 ul min^-1^ through a 100-μm needle connected to a pressure-controlled dispensing system (Smartsquirt, Automate Scientific). To avoid including responses due to mechanical stimulation of the cells, we applied aCSF through the needle and regarded this as the baseline before switching to pharmacological reagents. Pharmacological reagents used in this study are listed in the table below.

As a note, U73122 and U73343 stock solutions were prepared by first dissolving in chloroform to 10 mM. 250 μg aliquots were made in glass vials, and chloroform was evaporated away under a stream of Argon gas. The drugs were stored at −20°C for <1 week before re-dissolving in DMSO for use.

| Reagent | Description | Source | Working conc. | Application route |
| --- | --- | --- | --- | --- |
| Dynorphin A<sub>1-17</sub> | Endogenous κ agonist | Tocris 3195 | 1 µM | Locally for 30 sec |
| Nalfurafine | κ agonist | AdooQ BioScience A12579 | 1 µM (0.01% DMSO) | Locally for 30 sec |
| BRL-52537 | κ agonist | Abcam ab120669 | 10 µM (0.1% DMSO) | Locally for 30 sec |
| Nor-BNI | κ antagonist | Abcam ab120078 | 0.1 µM | Bath for 5-10 min |
| DIPPA | κ antagonist | Tocris 0794 | 1 µM (0.002% DMSO) | Bath for 5-10 min |
| Tetrodotoxin | Na<sub>v</sub> channel blocker | Cayman Chemical Company 14963 | 0.5 µM | Bath for 5-10 min |
| CNQX disodium | non-NMDA iGluR antagonist | Sigma-Aldrich C239 | 20 µM | Bath as a cocktail for 5-10 min |
| D-AP5 | NMDAR antagonist | Tocris 0106 | 50 µM |  |
| Picrotoxin | GABA<sub>A</sub> antagonist | Tocris 1128 | 100 µM (0.1% DMSO) |  |
| Strychnine HCl | GlyR antagonist | Sigma-Aldrich S8753 | 1 µM |  |
| Mecamylamine HCl | nAChR antagonist | Tocris 2843 | 100 µM |  |
| YM254890 | Gα<sub>q/11</sub> inhibitor | Tocris 7352 | 10 µM (0.1% DMSO) | Bath for 30 min |
| U73122 | PLC inhibitor | Tocris 1268 | 10 µM (0.2% DMSO) | Bath for 5-10 min |
| U73343 | Inactive analog of U73122 | Tocris 4133 | 10 µM (0.2% DMSO) | Bath for 5-10 min |
| Ivabradine HCl | HCN channel blocker | Sigma-Aldrich SML0281 | 10 µM | Bath for 5-10 min |
| Thapsigargin | SERCA ATPase inhibitor | Cayman Chemical Company 10522 | 4 µM (0.01% DMSO) | Bath for 10-15 min |
| Chelerythrine Cl | PKC inhibitor | Tocris 1330 | 10 µM | Bath for 30-40 min |
| Nifedipine | L-type Ca<sub>v</sub> channel blocker | Alomone Labs N-120 | 10 µM (0.01% DMSO) | Bath for 5-15 min |
| ω-Conotoxin GVIA | N-type Ca<sub>v</sub> channel blocker | Alomone Labs C-300 | 0.3 µM | Bath for 5-15 min |
| ω-Agatoxin IVA | P/Q-type Ca<sub>v</sub> channel blocker | Alomone Labs STA-500 | 0.3 µM | Bath for 10-15 min |
| SNX482 | R-type Ca<sub>v</sub> channel blocker | Tocris 2945 | 0.1 µM | Bath for 5-10 min |
| NNC 55-0396 | T-type Ca<sub>v</sub> channel blocker | Cayman Chemical Company 17216 | 25 µM | Bath for 5-15 min |

To investigate the role of external Ca^2+^, Ca^2+^-free solution (124 mM NaCl, 2.5 mM KCl, 4 mM MgSO_4_, 1.2 mM NaH_2_PO_4_, 24 mM NaHCO_3_, 5 mM HEPES, 1 mM EGTA, 12.5 mM Glucose, pH 7.4) was bath applied. At the end of each imaging session, high K^+^ solution (2.5 mM NaCl, 124 mM KCl, 2 mM MgSO4, 2 mM CaCl_2_, 1.2 mM NaH_2_PO_4_, 24 mM NaHCO_3_, 5 mM HEPES, 12.5 mM Glucose, pH 7.4) was squirted onto the slice to reveal all responsive neurons within the imaging field.

Fluorescence images were imported into the Fiji software (NIH), and regions of interest (ROIs) were drawn around labelled CSF-cNs. ΔF/F traces were extracted for each ROI over three time periods: (1) 30 sec immediately before DynA application, (2) 30 sec immediately following DynA application, and (3) 10 sec at the peak of the high K^+^ response. Since the response of CSF-cNs to DynA is typically composed of multiple transient spikes, we computed the area under ΔF/F traces for comparisons instead of using a single peak amplitude value. To account for the basal spontaneous activity of CSF-cNs and the difference in fluorescence due to the variable depths of the cells within the tissue slices, we subtracted away the baseline value and further normalized to the high K^+^ response. In other words, the DynA response is reported as

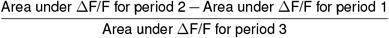

Slices that showed substantial movements during imaging or abnormal high K^+^ response were not analyzed. Cells with strong basal fluorescence (indicative of poor health) or large or extremely frequent spontaneous fluorescence fluctuations were also excluded from analyses due to difficulty in defining the baseline. ΔF/F images were generated by using the “F div F0” function in the T-functions plugin in Fiji.

### Patch-clamp recording

Spinal cord slices were prepared from *Tg(PKD2L1-Cre);Polr2a*^*GCaMP5G-tdTomato*^ mice by using the vibratome as above except at a thickness of 150 μm for better visualization under epifluorescence microscope (BX50WI, Olympus). During recording, slices were bath perfused with room-temperature aCSF bubbled with 95% O_2_/5% CO_2_ at a rate of ∼ 70 μl min^-1^. Patch electrodes (3-5 MΩ) were pulled from borosilicate capillaries (BF-150-110-10, Sutter Instrument) and filled with internal solution (10 mM NaCl, 135 mM KCl, 2 mM MgCl_2_, 1 mM CaCl_2_, 10 mM HEPES, 5 mM EGTA, 10 mM Phosphocreatine-Tris, 4 mM Mg-ATP, pH 7.4 with KOH). Under brightfield, the ependymal region was located at 4× magnification and CSF-cNs were identified at 40× by their tdTomato signal. We targeted only cells that were exposed or beneath a single cell layer. After the whole-cell patch-clamp configuration was established, voltage steps were given to confirm excitability of the cell. Recording was made by using the Multiclamp 700A amplifier (Molecular Devices) connected to a digitizer (Digidata 1550B, Molecular Devices) and was sampled at 10 kHz. For voltage-clamp recording, a low pass filter of 3 kHz (software single-pole RC filter) was applied. Cells were held at −80 mV without correcting for liquid-junction potential. As soon as a stable baseline was achieved, we locally perfused aCSF with a 100-μm Smartsquirt needle placed close to the ependymal region, followed by DYNA (1 μM), at a rate of ∼50 μl min^-1^. In experiments involving κ antagonist, 0.1 μM Nor-BNI was included in the bath. For current-clamp recording, low pass filter was set at 10 kHz and liquid-junction potential was corrected. When necessary, cells were brought to about −60 mV by current injection before DYNA application. Drugs were applied as above.

For analysis, we selected recordings that were made at an access resistance <25 MΩ and showed no drifting nor excessive noise. To produce an amplitude histogram, traces were further low-pass filtered digitally at 1 kHz (8-pole Bessel), and single channel opening events were identified by the Single-Channel Search function in the Clampfit software (v11.2.2.17, Molecular Devices) with open levels set at −4 pA and  −12 pA. The Gaussian function was fitted by using the Prism software (GraphPad). To study the effect of DYNA on open probability, the original 3 kHz filtering was maintained and single channel opening events were searched over a 30-sec stretch of recording with a single open level at  −12 pA in Clampfit and then confirmed manually by eye. Postsynaptic events were distinguished based on kinetics over the same period. Open probability was calculated by dividing the total dwell time by the recording duration. To calculate the firing rate, spikes were searched over a 30-sec stretch of recording by setting an amplitude threshold of 25 mV in Clampfit. Events were confirmed manually and firing rate was computed by dividing the number of spikes by the recording duration.

### Drug administration

Uninjured mice received daily injections of the κ opioid receptor agonist Nalfurafine (i.p. injected at a concentration of 4 μg/ml in sterile 0.9% saline with 0.1% DMSO to give a final dose of 20 μg/kg body weight, once/day, AdooQ Bioscience, A12579) or antagonist Nor-BNI (i.p. injected at a concentration of 2 mg/ml in sterile 0.9% saline to give a final dose of 10 mg/kg body weight, once/day, Abcam, ab120078) or the respective vehicle control over a course of 8 days. Starting from day 5 of the treatment, 50 mg/kg EdU (at a concentration of 10 mg/ml in sterile 0.9% saline, Cayman Chemical Company, 20518) was additionally i.p. injected once everyday for 7 days. Mice were sacrificed on the day after the EdU injections ended. All injections were done at roughly the same time of the day (∼5 PM) for consistency.

In spinal cord injury experiments, mice received the same 8-day agonist/antagonist treatment as above starting from the afternoon of the day of injury. EdU injections began on the following day and lasted for 7 days. Mice were sacrificed on the next day. For long-term drug administration, Nalfurafine powder was mixed with a matrix comprising cholesterol, cellulose, lactose, phosphates and stearates. The mixture was made into a pellet (0.027 mg Nalfurafine per pellet, 60-day release rate, Innovative Research of America). A placebo pellet containing only the matrix was used as a control. Five days before dorsal hemisection of the spinal cord, a pellet was implanted subcutaneously at the neck between the ears under isoflurane anesthesia, and the wound was closed with sutures and wound clips.

### Spinal cord hemisection

Under isoflurane anesthesia, hairs were removed from the dorsal surface of the mouse, and the surgical site was disinfected. An incision was made along the dorsal spine and the skin was held back by a retractor to expose the vertebrae. The muscles covering one of the lower thoracic or lumbar vertebral disks was cut in a manner perpendicular to the disk space. Laminectomy was then performed by using small vannas scissors to cut through the spinous processes and the lateral sides of the vertebral lamina. After lifting off the dorsal aspect of the vertebrae, a pair of microscissors was inserted into the exposed spinal cord segment at a depth of ∼0.8 mm to remove the dorsal column and dorsal horns. The muscle layer was closed by suturing and the skin was stapled. The mouse was supplemented with Lactated Ringer’s with 5% Dextrose (Baxter Healthcare Corporation, NDC 0338-0125-03) to regain homeostasis and was allowed to recover on a heating pad. Buprenorphine (Par Pharmaceutical, NDC 42023 −179-05, 0.1 mg/kg) and Meloxicam (Pivetal, NDC 46066-937 −13, 10 mg/kg) were administered i.p. before and after the surgery for analgesia. Sham-operated mice underwent the same procedure except that no spinal segment was removed. All surgeries were performed in the morning so that drug treatment could begin in the same afternoon.

During the first week after surgery and whenever necessary thereafter, we manually expressed the bladder of the animals by using the Crede maneuver twice daily. Both EAE and BCS scores were recorded daily. None of the operated mice used in this study had a >15% drop in body weight.

### Data analysis and statistics

Unless otherwise specified, collective data are reported as mean ± standard deviation (SD). Statistical analyses were done by using the Prism software (GraphPad), and data are reported following the ARRIVE guidelines. Statistical comparisons were done by using the Welch’s t test, Dunnett test or ANOVA as specified in the figure legends. Significance was determined by the computed p-value. *p ≤ 0.05; **p ≤ 0.01; ***p ≤ 0.001; ****p ≤ 0.0001; ns: not significant.

## Data availability

All data generated or analyzed during this study are included in the manuscript (and its supplementary information data files and the source data files). The sequencing results will be deposited at NCBI. Source data are provided with this paper. Reagents and codes used are available upon request.

## Inclusion and ethics statement

We support inclusive, diverse, and equitable conduct of research.

## Acknowledgements

We thank Dr. Charles Zuker (Columbia University) for providing the *Tg(PKD2L1-Cre)* mouse line, Dr. Nicholas Ingolia (University of California, Berkeley) for providing computational resources for sequencing analyses and Ms. Jeannie Poblete for technical support. We also thank Drs. Roger Nicoll, Michael Beattie, Jacqueline Bresnahan, Allan Basbaum, Joao Braz, Markus Delling, Nicholas Bellono, Zheng Jiang, Kevin Yackle, and all current members of the Julius lab for discussion and critical reading of the manuscript. We appreciate the support from staff in UCSF’s core facilities, including the Laboratory for Cell Analysis (Dr. Sarah Elmes, NIH Cancer Center Support Grant P30CA082103), Center for Advanced Light Microscopy (Drs. DeLaine Larsen, Kari Herrington and So Yeon Kim, S10 Shared Instrumentation Grant 1S10OD017993-01A1 for Nikon CSU-W1 spinning disk confocal microscope), Center for Advanced Technology (Drs. Eric Chow and Delsy Martinez) as well as the Mouse Microsurgery Core (Drs. Mark Looney and Longhui Qiu, financial support from UCSF Bakar ImmunoX Initiative). This work was supported by a Howard Hughes Medical Institute Hanna Gray Fellowship and a Croucher Fellowship for Postdoctoral Research (to WWSY), a Damon Runyon Cancer Research Foundation Fellowship (DRG-[2387-30] to KKT), the UCSF Program for Breakthrough Biomedical Research: New Frontier Research Award (to DJ) and NIH grants (R01EY030138 to XD and R35 NS105038 to DJ).

## Supplementary data

**Supplementary Fig. 1.**
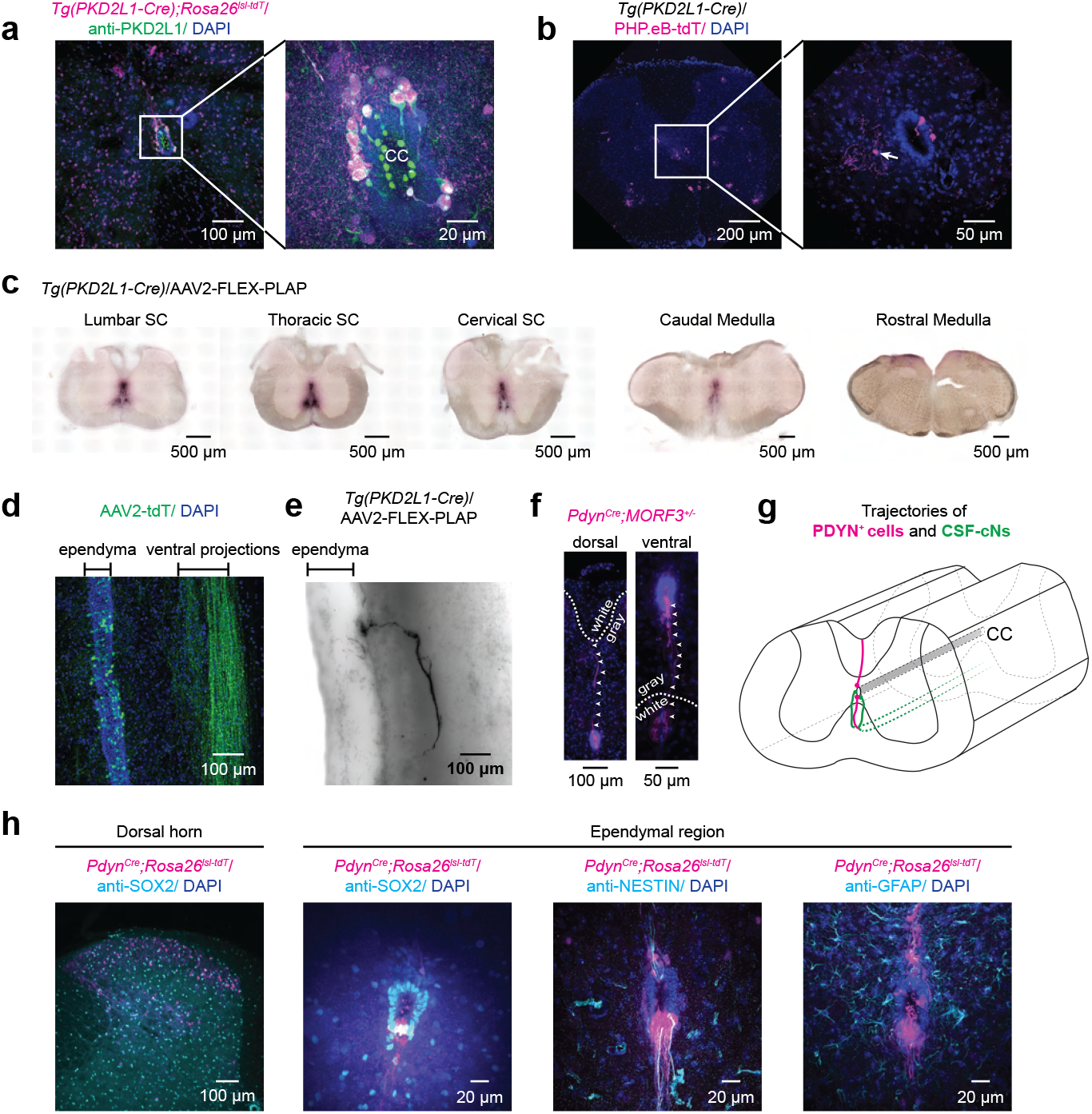
Characterization of CSF-cNs and PDYN^+^ cells in the spinal cord. **a**, Labeling of spinal cord cells by the *Tg(PKD2L1-Cre);Rosa26*^*LSL-tdTomato*^ mouse line. The tdTomato reporter (magenta) was expressed in many other spinal cord cells in addition to the CSF-cNs (magnified on right), which were identified by their strong immunostaining signal for PKD2L1 (green), particularly in their bulbous projections within the central canal (CC). **b**, Labeling of spinal cord cells by the *Tg(PKD2L1-Cre)* mouse line when tdTomato reporter was delivered by i.c.v. AAV injection after adulthood. The PHP.eB serotype was used because it crosses the ependymal barrier. Cells outside of the ependymal region were labelled (e.g., arrow in magnified view on right). **c**, Projection pattern of CSF-cNs at different levels of the spinal cord. Coronal spinal cord sections from a mouse that had received i.c.v. injection of an AAV carrying the PLAP transgene. Sections were stained with NBT/BCIP to reveal the ventral projections of CSF-cNs. **d**, Longitudinal projection of CSF-cNs. Sagittal section of the spinal cord showing the nerve bundle formed by CSF-cNs’ projections that travelled rostrocaudually within the ventral white matter. Cell bodies of CSF-cNs are found in the ependyma. **e**, Morphology of a single PLAP-labelled CSF-cN. Spinal cord was cleared by the iDisco method (46) after NBT/BCIP staining. **f**, Morphology of *Pdyn*^*+*^ cells. Arrowheads trace the long processes of sparsely labelled *Pdyn*^*+*^ cells situated at the dorsal or ventral pole of the ependyma. Dotted lines mark the boundary between gray and white matters. **g**, Schematic depicting the projection pattern of CSF-cNs (green) and PDYN^+^ cells (magenta). **h**, Spinal cord sections from *Pdyn*^*Cre*^*;Rosa26*^*tdTomato*^ mice immunostained with antibodies against known markers (cyan) for various subsets of ependymal cells. SOX2 is expressed in PDYN^+^ cells (magenta) in the ependymal region but not those in the dorsal horn. Ependymal PDYN^+^ cells are also positive for NESTIN but not GFAP.

**Supplementary Fig. 2.**
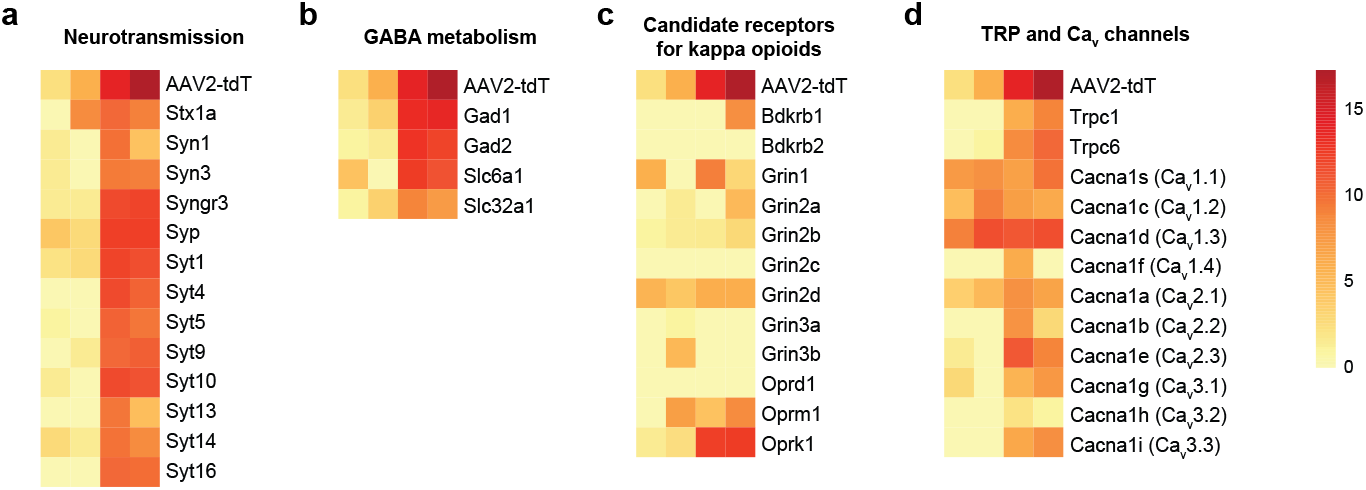
Expression of transcripts of interest in preparations of acutely dissociated and FACS-enriched CSF-cNs. CSF-cNs were labelled by tdTomato via i.c.v. AAV injection and were fluorescently sorted for bulk mRNA sequencing. Columns represent 4 separate preparations with increasing level of enrichment in CSF-cNs, as reflected by the expression level of *tdTomato* transcript. Color scale is based on median-of-ratios calculation by DESeq2. **a**, Genes related to neurotransmission. **b**, Genes typically involved in GABA metabolism. **c**, Genes for receptors proposed to be sensitive to κ opioid ligands. **d**, TRP and Ca_v_ channel genes.

**Supplementary Fig. 3.**
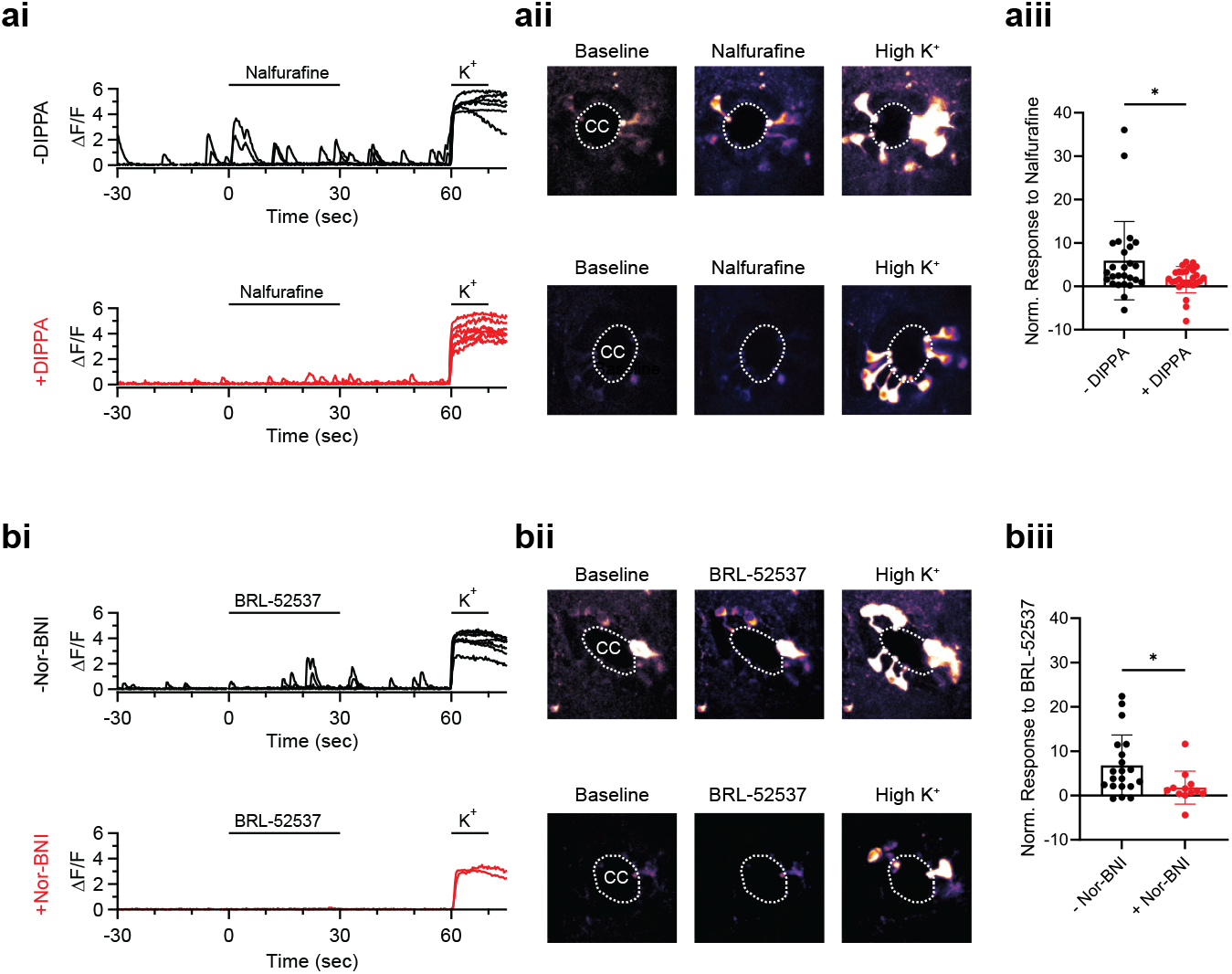
Ca^2+^ responses of GCaMP5G-expressing CSF-cNs to OPRK1 agonists. **ai**, Example ΔF/F traces showing the responses of CSF-cNs to local application of the κ agonist, Nalfurafine, in the absence (black) or presence (red) of the antagonist, DIPPA, in the bath. Local application of a high K^+^ solution was used to reveal all responsive neurons. Each trace is from a single cell. **aii**, ΔF/F images for the spinal cord slices in **ai**. Images are temporal averages over 10 sec of baseline or for the duration of the stimuli. CC: central canal. **aiii**, Collective data comparing CSF-cNs’ responses to Nalfurafine in the absence (black) and presence (red) of DIPPA. Each dot shows the integral DYNA response of a single cell normalized to the high-K^+^ response. Welch’s t test, *p ≤ 0.05, n = 26 and 27. **b**, Same as a except that BRL-52537 was used as the agonist and Nor-BNI as the antagonist. n = 20 and 12.

**Supplementary Fig. 4.**
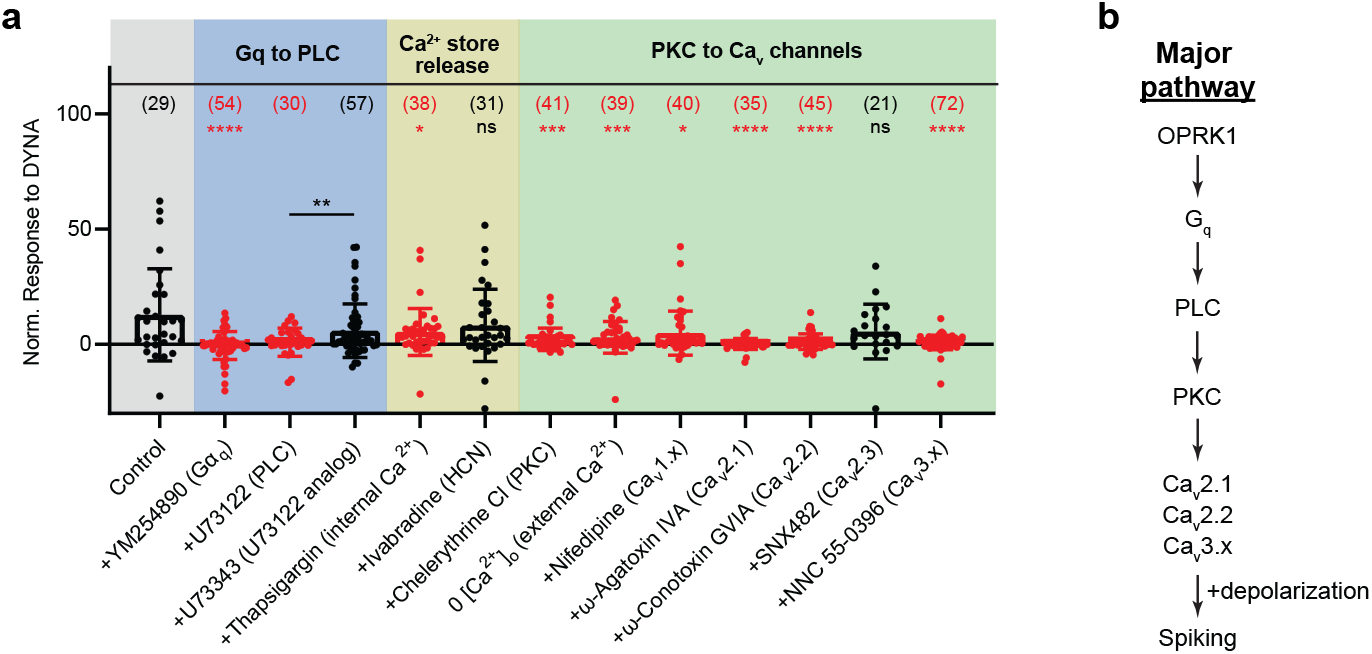
Pharmacological experiments to delineate the downstream pathway of OPRK1 signaling. Normalized integral DYNA response as in **Fig. 2c**, but in the presence of various inhibitors or in different ionic conditions. Molecular targets of the drugs are indicated in brackets. Routes of drug application are detailed in Methods. Numbers of cells used are in brackets above bars. Post-hoc Dunnett tests for comparison with control, which is same as the – Nor-BNI condition in **Fig. 2biii** and **Fig. 2c**, *p ≤ 0.05; ***p ≤ 0.001; ****p ≤ 0.0001 (highlighted in red); ns: not significant (black). The effect of the PLC inhibitor, U73122, was compared to that of its inactive analog, U73343, prepared in the same solvent (see Methods); Welch’s t test. **b**, Proposed signaling pathway downstream of OPRK1 in CSF-cNs.

**Supplementary Fig. 5.**
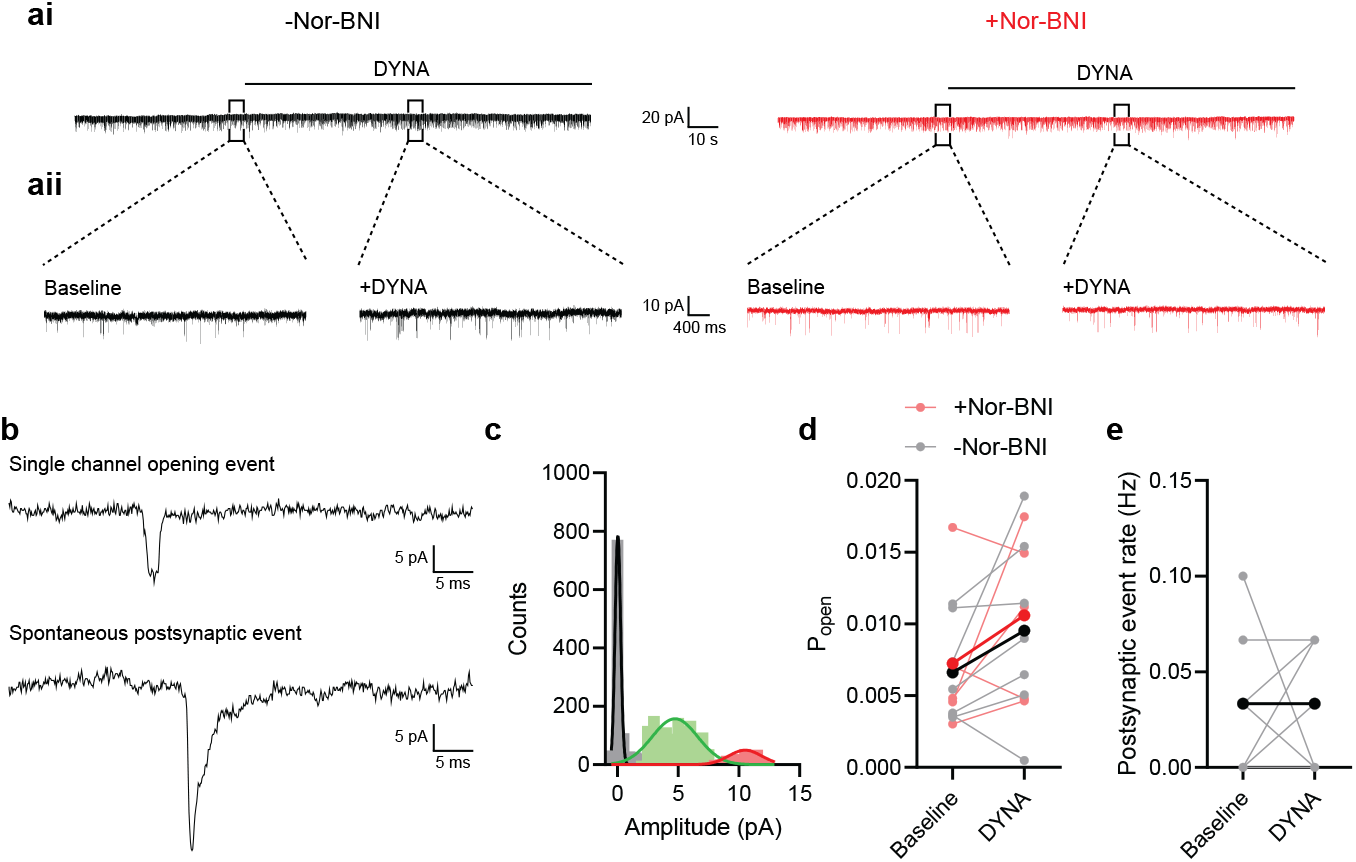
Voltage-clamp recording on CSF-cNs during DYNA application. **ai**, Recording traces from two representative cells voltage-clamped at −80 mV in the absence or presence of the κ antagonist, Nor-BNI, in bath. No macroscopic current was observed during local DYNA application (line above trace). **aii**, Expanded view of the boxed regions of traces in **ai**, showing single channel openings at baseline or during DYNA application. **b**, Example of a single channel opening event and a spontaneous postsynaptic event to show the clear distinction between the two waveforms. **c**, Amplitude histogram of spontaneous single channel opening events detected at baseline. Two peaks at amplitude ∼5 pA and ∼11 pA were detected. The ∼11 pA events resembled those described in earlier reports (11, 15), which were shown to come from PKD2L1 channels (15). **d**, Open probability of the ∼11 pA channel before and after DYNA application with or without Nor-BNI in bath. Each pair of light-colored dots is from a single cell. Group averages are in dark colors. No statistical significance detected with two-way ANOVA across DYNA or antagonist condition (n = 6 and 6). We did not analyze the ∼5 pA events because they were not clearly discernable from background noise. **e**, Rate of spontaneous postsynaptic event before and during DYNA application in normal aCSF bath. Each pair of light-colored dots is from a single cell. Group averages are in dark colors. No statistical significance detected with paired t-test (n = 7 and 7).

**Supplementary Fig. 6.**
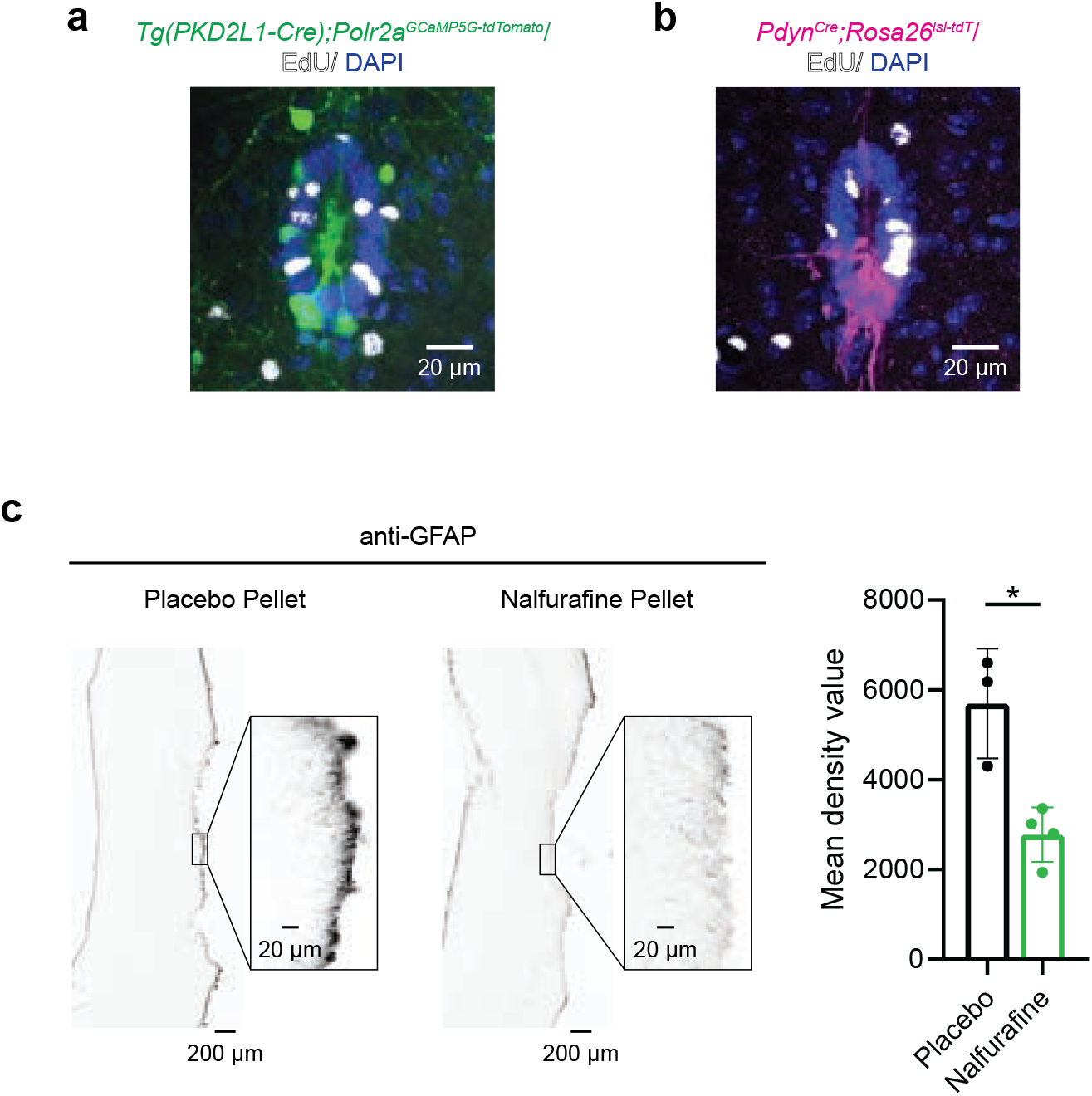
CSF-cNs, *Pdyn*^*+*^ cells and scar formation in injured spinal cords. Following dorsal hemisection, no EdU signal (white) was detected in (**a**) CSF-cNs labelled in the *Tg(PKD2L1-Cre)* mouse line and (**b**) *Pdyn*^*+*^ cells labelled in the *Pdyn*^*Cre*^ line on Day 9. EdU injection scheme as in **Fig. 4b. c**, Representative sagittal images of injured spinal cords harvested from placebo- or Nalfurafine-treated mice 5 weeks after dorsal hemisection. Sections were immunostained with GFAP antibody to label glial scar. Intensity of immunosignal along the perimeter of the lesion site was quantified on right. Welch’s t-test, *p ≤ 0.05, n = 3 and 4.

## References

1. Silver, J., and Miller, J. H. (2004) Regeneration beyond the glial scar. Nature Reviews Neuroscience 5, 146–156.

2. Tran, A. P., Warren, P. M., and Silver, J. (2021) New insights into glial scar formation after spinal cord injury. Cell and Tissue Research 387, 319–336.

3. Johansson, C. B., Momma, S., Clarke, D. L., Risling, M., Lendahl, U., and Frisén, J. (1999) Identification of a neural stem cell in the adult mammalian central nervous system. Cell 96, 25–34.

4. Meletis, K., Barnabé-Heider, F., Carlén, M., Evergren, E., Tomilin, N., Shupliakov, O., and Frisén, J. (2008) Spinal cord injury reveals multilineage differentiation of ependymal cells. PLOS Biology 6, e182.

5. Sabelström, H., Stenudd, M., Réu, P., Dias, D. O., Elfineh, M., Zdunek, S., Damberg, P., Göritz, C., and Frisén, J. (2013) Resident neural stem cells restrict tissue damage and neuronal loss after spinal cord injury in mice. Science (New York, N.Y.) 342, 637–640.

6. Barnabé-Heider, F., Göritz, C., Sabelström, H., Takebayashi, H., Pfrieger, F. W., Meletis, K., and Frisén, J. (2010) Origin of new glial cells in intact and injured adult spinal cord. Cell Stem Cell 7, 470–482.

7. Lacroix, S., Hamilton, L. K., Vaugeois, A., Beaudoin, S., Breault-Dugas, C., Pineau, I., Lévesque, S. A., Grégoire, C. A., and Fernandes, K. J. (2014) Central canal ependymal cells proliferate extensively in response to traumatic spinal cord injury but not demyelinating lesions. PLOS ONE 9, e85916.

8. New, L. E., Yanagawa, Y., McConkey, G. A., Deuchars, J., and Deuchars, S. A. (2023) GABAergic regulation of cell proliferation within the adult mouse spinal cord. Neuropharmacology 223, 109326.

9. Vigh, B., Vigh-Teichmann, I., Manzano e Silva, M. J., and van den Pol, A. N. (1983) Cerebrospinal fluid-contacting neurons of the central canal and terminal ventricle in various vertebrates. Cell and Tissue Research 231, 615–621.

10. Huang, A. L., Chen, X., Hoon, M. A., Chandrashekar, J., Guo, W., Tränkner, D., Ryba, N. J. P., and Zuker, C. S. (2006) The cells and logic for mammalian sour taste detection. Nature 442, 934–938.

11. Orts-Del’immagine, A., Wanaverbecq, N., Tardivel, C., Tillement, V., Dallaporta, M., and Trouslard, J. (2012) Properties of subependymal cerebrospinal fluid contacting neurones in the dorsal vagal complex of the mouse brainstem. J Physiol 590, 3719–3741.

12. Prendergast, A. E. et al. (2023) CSF-contacting neurons respond to Streptococcus pneumoniae and promote host survival during central nervous system infection. Current Biology

13. Böhm, U. L., Prendergast, A., Djenoune, L., Nunes Figueiredo, S., Gomez, J., Stokes, C., Kaiser, S., Suster, M., Kawakami, K., Charpentier, M., Concordet, J.-P., Rio, J.-P., Del Bene, F., and Wyart, C. (2016) CSF-contacting neurons regulate locomotion by relaying mechanical stimuli to spinal circuits. Nature Communications 7, 1–8.

14. Sternberg, J. R., Prendergast, A. E., Brosse, L., Cantaut-Belarif, Y., Thouvenin, O., Orts-Del’Immagine, A., Castillo, L., Djenoune, L., Kurisu, S., McDearmid, J. R., Bardet, P.-L., Boccara, C., Okamoto, H., Delmas, P., and Wyart, C. (2018) Pkd2l1 is required for mechanoception in cerebrospinal fluid-contacting neurons and maintenance of spine curvature. Nature Communications 9, 3804.

15. Orts-Del’Immagine, A., Seddik, R., Tell, F., Airault, C., Er-Raoui, G., Najimi, M., Trouslard, J., and Wanaverbecq, N. (2016) A single polycystic kidney disease 2-like 1 channel opening acts as a spike generator in cerebrospinal fluid-contacting neurons of adult mouse brainstem. Neuropharmacology 101, 549–565.

16. Johnson, E., Clark, M., Oncul, M., Pantiru, A., MacLean, C., Deuchars, J., Deuchars, S. A., and Johnston, J. (2023) Graded spikes differentially signal neurotransmitter input in cerebrospinal fluid contacting neurons of the mouse spinal cord. iScience 26, 105914.

17. Gerstmann, K., Jurčić, N., Blasco, E., Kunz, S., de Almeida Sassi, F., Wanaverbecq, N., and Zampieri, N. (2022) The role of intraspinal sensory neurons in the control of quadrupedal locomotion. Current Biology 32, 2442–2453.e4.

18. Djenoune, L., Desban, L., Gomez, J., Sternberg, J. R., Prendergast, A., Langui, D., Quan, F. B., Marnas, H., Auer, T. O., Rio, J. P., Del Bene, F., Bardet, P. L., and Wyart, C. (2017) The dual developmental origin of spinal cerebrospinal fluid-contacting neurons gives rise to distinct functional subtypes. Scientific Reports 7, 1–14.

19. Nakamura, Y. et al. (2023) Cerebrospinal fluid-contacting neuron tracing reveals structural and functional connectivity for locomotion in the mouse spinal cord. eLife 12.

20. Stoeckel, M.-E., Uhl-Bronner, S., Hugel, S., Veinante, P., Klein, M.-J., Mutterer, J., Freund-Mercier, M.-J., and Schlichter, R. (2003) Cerebrospinal fluid-contacting neurons in the rat spinal cord, a γ-aminobutyric acidergic system expressing the P2X2 subunit of purinergic receptors, PSA-NCAM, and GAP-43 immunoreactivities: light and electron microscopic study. The Journal of Comparative Neurology 457, 159–174.

21. Chavkin, C. (2013) Dynorphin–still an extraordinarily potent opioid peptide. Molecular Pharmacology 83, 729–736.

22. Khachaturian, H., Watson, S. J., Lewis, M. E., Coy, D., Goldstein, A., and Akil, H. (1982) Dynorphin immunocytochemistry in the rat central nervous system. Peptides 3, 941–954.

23. Veldman, M. B. et al. (2020) Brainwide genetic sparse cell labeling to illuminate the morphology of neurons and glia with Cre-dependent MORF mice. Neuron 108, 111–127.e6.

24. Furube, E., Ishii, H., Nambu, Y., Kurganov, E., Nagaoka, S., Morita, M., and Miyata, S. (2020) Neural stem cell phenotype of tanycyte-like ependymal cells in the circumventricular organs and central canal of adult mouse brain. Scientific Reports 2020 10:1 10, 1–15.

25. Brust, T. F. (2022) Biased ligands at the kappa opioid receptor: fine-tuning receptor pharmacology. Handbook of Experimental Pharmacology 271, 115–135.

26. Bruchas, M. R., and Chavkin, C. (2010) Kinase cascades and ligand-directed signaling at the kappa opioid receptor. Psychopharmacology 210, 137–147.

27. Eriksson, P. S., Nilsson, M., Wågberg, M., Hansson, E., and Rönnbäck, L. (1993) Kappa-opioid receptors on astrocytes stimulate l-type Ca2+ channels. Neuroscience 54, 401–407.

28. Gurwell, J. A., Duncan, M. J., Maderspach, K., Stiene-Martin, A., Elde, R. P., and Hauser, K. F. (1996) κ-Opioid receptor expression defines a phenotypically distinct subpopulation of astroglia: relationship to Ca2+ mobilization, development, and the antiproliferative effect of opioids. Brain Research 737, 175–187.

29. Pan, Z. Z. (2003) κ-Opioid receptor-mediated enhancement of the hyperpolarization-activated current (Ih) through mobilization of intracellular calcium in rat nucleus raphe magnus. The Journal of Physiology 548, 765–775.

30. Lai, J., Luo, M. C., Chen, Q., Ma, S., Gardell, L. R., Ossipov, M. H., and Porreca, F. (2006) Dynorphin A activates bradykinin receptors to maintain neuropathic pain. Nature Neuroscience 2006 9:12 9, 1534–1540.

31. Laughlin, T. M., Vanderah, T. W., Lashbrook, J., Nichols, M. L., Ossipov, M., Porreca, F., and Wilcox, G. L. (1997) Spinally administered dynorphin A produces long-lasting allodynia: involvement of NMDA but not opioid receptors. Pain 72, 253–260.

32. Bakshi, R., and Faden, A. I. (1990) Competitive and non-competitive NMDA antagonists limit dynorphin A-induced rat hindlimb paralysis. Brain Research 507, 1–5.

33. Zhang, S., Tong, Y., Tian, M., Dehaven, R. N., Cortesburgos, L., Mansson, E., Simonin, F., Kieffer, B., and Yu, L. (1998) Dynorphin A as a potential endogenous ligand for four members of the opioid receptor gene family. The Journal of pharmacology and experimental therapeutics 286, 136–141.

34. Corns, L. F., Atkinson, L., Daniel, J., Edwards, I. J., New, L., Deuchars, J., and Deuchars, S. A. (2015) Cholinergic enhancement of cell proliferation in the postnatal neurogenic niche of the mammalian spinal cord. Stem cells (Dayton, Ohio) 33, 2864–2876.

35. Hussein, S. A. (2015) Functional Characterization of the TRP-Type Channel PKD2L1.

36. Felix, R. (2008) Molecular regulation of voltage-gated Ca2+ channels. Journal of Receptors and Signal Transduction 25, 57–71.

37. Barber, R. P., Vaughn, J. E., and Roberts, E. (1982) The cytoarchitecture of GABAergic neurons in rat spinal cord. Brain Research 238, 305–328.

38. Djenoune, L., Khabou, H., Joubert, F., Quan, F. B., Figueiredo, S. N., Bodineau, L., Del Bene, F., Burcklé, C., Tostivint, H., and Wyart, C. (2014) Investigation of spinal cerebrospinal fluid-contacting neurons expressing PKD2L1: Evidence for a conserved system from fish to primates. Frontiers in Neuroanatomy 8, 26.

39. Ren, Y., Ao, Y., O’Shea, T. M., Burda, J. E., Bernstein, A. M., Brumm, A. J., Muthusamy, N., Ghashghaei, H. T., Carmichael, S. T., Cheng, L., and Sofroniew, M. V. (2017) Ependymal cell contribution to scar formation after spinal cord injury is minimal, local and dependent on direct ependymal injury. Scientific Reports 7, 1–16.

40. Corns, L. F., Deuchars, J., and Deuchars, S. A. (2013) GABAergic responses of mammalian ependymal cells in the central canal neurogenic niche of the postnatal spinal cord. Neuroscience Letters 553, 57–62.

41. Snyder, L. M. et al. (2018) Kappa opioid receptor distribution and function in primary afferents. Neuron 99, 1274–1288.e6.

42. Inui, S. (2015) Nalfurafine hydrochloride to treat pruritus: a review. Clinical, Cosmetic and Investigational Dermatology 8, 249.

43. Madisen, L., Zwingman, T. A., Sunkin, S. M., Oh, S. W., Zariwala, H. A., Gu, H., Ng, L. L., Palmiter, R. D., Hawrylycz, M. J., Jones, A. R., Lein, E. S., and Zeng, H. (2010) A robust and high-throughput Cre reporting and characterization system for the whole mouse brain. Nature neuroscience 13, 133–140.

44. Gee, J. M. et al. (2014) Imaging activity in neurons and glia with a Polr2a-based and Cre-dependent GCaMP5G-IRES-tdTomato reporter mouse. Neuron 83, 1058–1072.

45. Buch, T., Heppner, F. L., Tertilt, C., Heinen, T. J. A. J., Kremer, M., Wunderlich, F. T., Jung, S., and Waisman, A. (2005) A Cre-inducible diphtheria toxin receptor mediates cell lineage ablation after toxin administration. Nature Methods 2, 419–426.

46. Renier, N., Wu, Z., Simon, D. J., Yang, J., Ariel, P., and Tessier-Lavigne, M. (2014) iDISCO: a simple, rapid method to immunolabel large tissue samples for volume imaging. Cell 159, 896–910.

